# Duckweeds as Multiplexable Plant Models for Exploring Abiotic Stress Responses

**DOI:** 10.64898/2026.09.03.749270

**Authors:** Bryan Ramirez-Corona, Santiago Cadena, Kerry Bubb, Christine Queitsch, Josh T. Cuperus

## Abstract

In nature, plants experience complex combinations of environmental stresses. Prior studies using *Arabidopsis thaliana* show that transcriptional responses to combinatorial stress are unique and cannot be predicted from responses to individual treatments, making it imperative to study combinatorial stress in the monocotyledonous cereals that comprise the majority of our food supply. However, crop species are not easily amenable for such studies because of their large size, long generation time, and organismal complexity. In contrast, the monocotyledonous aquatic duckweeds are small, grow rapidly, and show reduced organismal and genomic complexity. Here, we explore the physiological responses to combinatorial stress in three duckweed species, *S. polyrhiza*, *L. minor*, and *W. australiana* and examine their potential to serve as multiplexable models. Focusing on *S. polyrhiza,* the most complex and best annotated species, we investigated transcriptional response to individual and combinatorial stress treatments. While most individual treatments elicit few transcriptional changes, combinatorial stress elicited a non-additive transcriptional response unique to each combination. Exploring the duckweed regulatory landscape, we found that regulatory regions near genes that were differentially expressed in combinatorial stress in *S. polyrhiza* resemble environmentally responsive regulatory elements found in terrestrial plants. Examined duckweed species show numbers and genomic distributions of regulatory elements similar to those of *A. thaliana* and maize, with subtle effects of genome size but none of organismal complexity. Taken together, our results establish that duckweeds respond to combinatorial stress in a manner similar to terrestrial plants and can therefore serve as high-throughput, multiplexable models for studying its molecular underpinnings.

## Introduction

Plants provide the food that sustains our world and cumulatively comprise 80% of the earth’s biomass (Bar-On *et al*. 2018). In order to thrive, plants must contend with complex and dynamic combinations of both abiotic and biotic factors (Mittler 2006). The intricate mechanisms of environmental sensing, signal integration, and decision control that plants employ to interpret and respond to their complex environment are not fully understood (Mittler and Blumwald 2010). Faced with the ongoing loss of arable land and the rapid changes in climate patterns, we need a more sophisticated understanding of how plants, especially our monocotyledonous staple cereals, respond to combinatorial stress conditions (Pascual *et al*. 2022).

Historically, plant stress responses have been studied by exposing plants to single, well-defined stresses such as heat stress or herbivory. These studies have been invaluable in furthering our understanding of the pathways and genes involved in environmental responses. However, recent work has shown that individual stress treatments that alone have little effect can drastically reduce plant growth, productivity, and survival when applied in combination (Jiang and Huang 2001, Rizhsky *et al*. 2002, Zandalinas *et al*. 2021, Pascual *et al*. 2023, Sinha *et al*. 2024, Peláez-Vico *et al*. 2024, Sinha *et al*. 2025). Moreover, transcriptional responses to individual stress treatments do not predict transcriptional responses to combinatorial stress (Rizhsky *et al*. 2002, Rasmussen *et al*. 2013, Ristova *et al*. 2016, Zandalinas *et al*. 2021). These findings have motivated studies into the combinatorial stress response in crop plants (Pascual *et al*. 2023, Sinha *et al*. 2024, Peláez-Vico *et al*. 2024). However, to fully explore the vast combinatorial space of environmental stress responses and understand the underlying principles, plant models are needed that allow high throughput, multiplexed experiments.

One potential model for such experiments is *Spirodela polyrhiza*, a member of the *Lemnaceae* family, commonly known as duckweeds. This family of aquatic monocots comprises 36 species spanning five genera and contains both the smallest and the fastest-growing flowering plants (Acosta *et al*. 2021). The duckweeds’ small size, simplified body plan, and extremely rapid growth make them well-suited for studying combinatorial stress responses in high throughput and multiplexed (Di Stilio & Sinha 2024). *S. polyrhiza* has major plant tissues, such as a fused leaf stem-like structure, primary roots, meristem, vasculature, and flowers. The species’ aquatic lifestyle allows for easy maintenance and highly controlled treatment conditions. Thus far, there have been few reports of duckweed transcriptomes and regulatory landscapes, and high-quality duckweed genomes have only recently become available, with the *S. polyrhiza* genome being the most contiguous and best-annotated (Earnst *et al*. 2025).

To address whether *S. polyrhiza* is a suitable monocot model for studying combinatorial stress responses, we collected growth rate and survival data for this species in response to six abiotic treatments and their combinations. We also included two other commonly used duckweed species for comparison, *L. minor,* which offers existing phenotype data, and *W. australiana*, which has a simpler body plan than *S. polyrhiza*. The growth rate data informed our choice of treatment conditions for examining *S. polyrhiza* transcriptomes. The *S. polyrhiza* transcriptome data showed largely muted responses to individual stress treatments while combinations with temperature stress altered gene expression in a non-additive manner, leading us to conclude that duckweeds respond to combinatorial stress like terrestrial plants. Lastly, we used chromatin accessibility data in *S. polyrhiza* to find evidence of regulatory elements and trans-acting factors specific to combinatorial stress and compare chromatin accessibility across the three duckweed species, in addition to *A. thaliana* and maize. Our results demonstrate that duckweeds can serve as high throughput, multiplexable models for investigating the molecular underpinnings of complex stress responses in monocots.

## Results

### Determining duckweed growth rates in response to abiotic stress

To determine growth rates of *S. polyrhiza* 9509 in response to abiotic stress treatments, we challenged fronds with increasing concentrations of several compounds, increased growth temperature and low pH. We quantified growth by counting the total number of visible fronds (Material and Methods, Figure 1A and B). For comparison, we included two other, less complex and smaller duckweed species *Lemna minor* 7210 and *Wolffia australiana* 8730. The tested conditions included: temperature (TEMP), phosphate (PO4), nitrate (NO3), ammonium (NH4), salt (NACL), and low pH (HCL). We selected these conditions because duckweeds are often found in wastewater where they experience an overabundance of nutrients due to agricultural run-off (Acosta *et al*. 2021). High temperature and salt stress have broad relevance for all plants. As expected, increasing the relative stress levels generally resulted in decreased growth with the strongest stress treatments resulting in death (Figure 1C, Supplemental Figure 1A-B).

**Figure 1:**
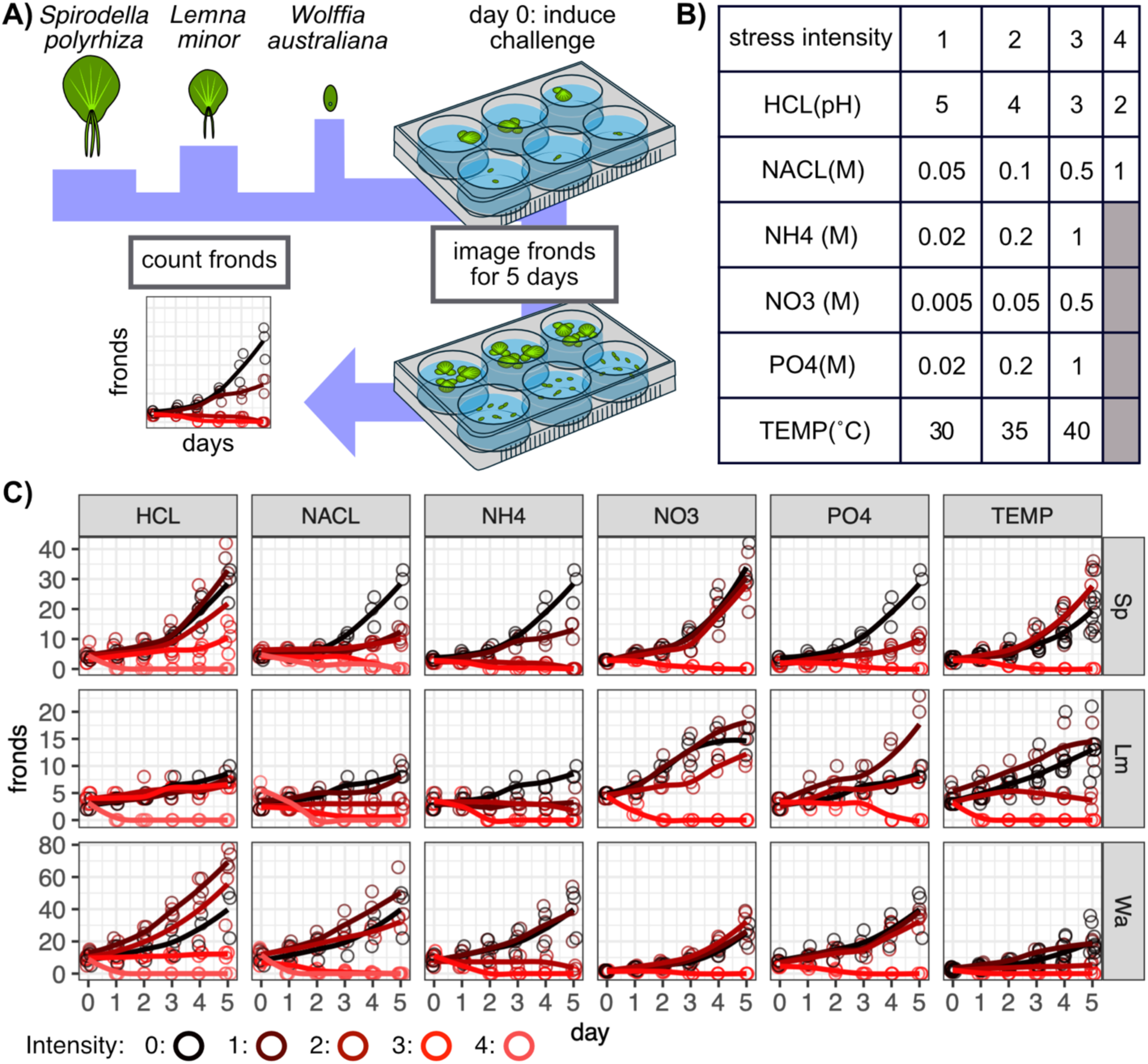
Duckweed species show distinct sensitivity profiles to abiotic stress. A) Schematic of experimental set up used to quantify growth in three duckweed species: *S. polyrhiza*, *L. minor*, and *W. australiana*. B) Treatment intensity (columns) for the tested abiotic stresses (rows). C) Growth curves for three duckweed species under varying degrees of stress intensity. Total live fronds (y-axis) were tracked over the course of five days (x-axis). Experimental conditions are colored by stress intensity with black denoting control and the brightest red denoting the highest tested intensity.

To compare growth rate effects among treatments, frond counts were log-transformed, and the slope was calculated for each species and condition (Supplemental Figure 1A). In control conditions, *L. minor* (m=0.1) showed the slowest growth with *S. polyrhiza* (m=0.18) and *W. australiana* (m=0.18) showing similar growth rates (Materials and Methods, Supplemental Figure 1B). *S. polyrhiza* appeared to have the highest resistance to temperature stress, showing growth reduction only at the highest temperature tested (40°C). In contrast; *L. minor* and *W. australiana* showed reduced growth at 30°C. All three species proved resilient to increases in phosphate and nitrate concentration, showing drastic growth reduction only at the highest concentrations tested. *S. polyrhiza* was particularly sensitive to salt and ammonium stress. *W. australiana* showed the highest resistance to low pH.

In *A. thaliana*, combinations of individual stress treatments drastically reduce survival (Rasmussen 2013, Zandalinas 2021). Thus, we hypothesized that combinatorial stress will similarly reduce survival in duckweeds. However, the aquatic lifestyle, minimal body plan, and reduced genomes of duckweeds might alter their response. We quantified survival of *S. polyrhiza* in response to combinations of two, four and five different stress treatments, including *L. minor* and *W. australiana* for comparison (Supplemental Figure 2A). Similar to results reported for terrestrial plants, increasing the complexity of stress treatments further reduced growth rate (Supplemental Figure 2B). This result suggests that duckweeds react to combinatorial stress like terrestrial plants.

### S. polyrhiza shows unique transcriptional responses to combinatorial stress

In *A. thaliana*, combinatorial stress results in unique transcriptional responses such that responses to individual stress treatments are not predictive of responses to combinatorial stress (Rasmussen 2013, Zandalinas 2021). We hypothesized that duckweeds might also show unique transcriptional responses to combinatorial stress. To test this hypothesis, we characterized both growth and transcriptomes in response to individual and combinatorial stress treatments in *S. polyrhiza*. Plants were either challenged with one of five media treatments or a high temperature stress or with combinations of a media and temperature stress. We intentionally combined mild media stress treatments with the temperature treatment to amplify possible unique growth and transcriptional responses to combinatorial stress.

Of the individual treatments, increased temperature resulted in the greatest reduction of growth rate, followed by salt stress (m_CONTROL_=0.16, m_TEMP_=0.044, m_NACL_=0.09, Figure 2A). The other treatments did not noticeably affect growth rates. The combinatorial stress treatments yielded reduced growth rates similar to those observed with temperature stress alone (m_TEMP+HCL_= 0.033, m_TEMP+NACL_= 0.021, m_TEMP+NH4_= 0.055, m_TEMP+PO4_= 0.044). The combination of nitrate treatment with temperature stress showed the greatest reduction in growth, slower than expected for the sum of both individual treatments (m_TEMP+NO3_=-0.041). However, frond growth responses to combinatorial treatments overall did not significantly deviate from additive expectations (Supplemental Table 1).

**Figure 2:**
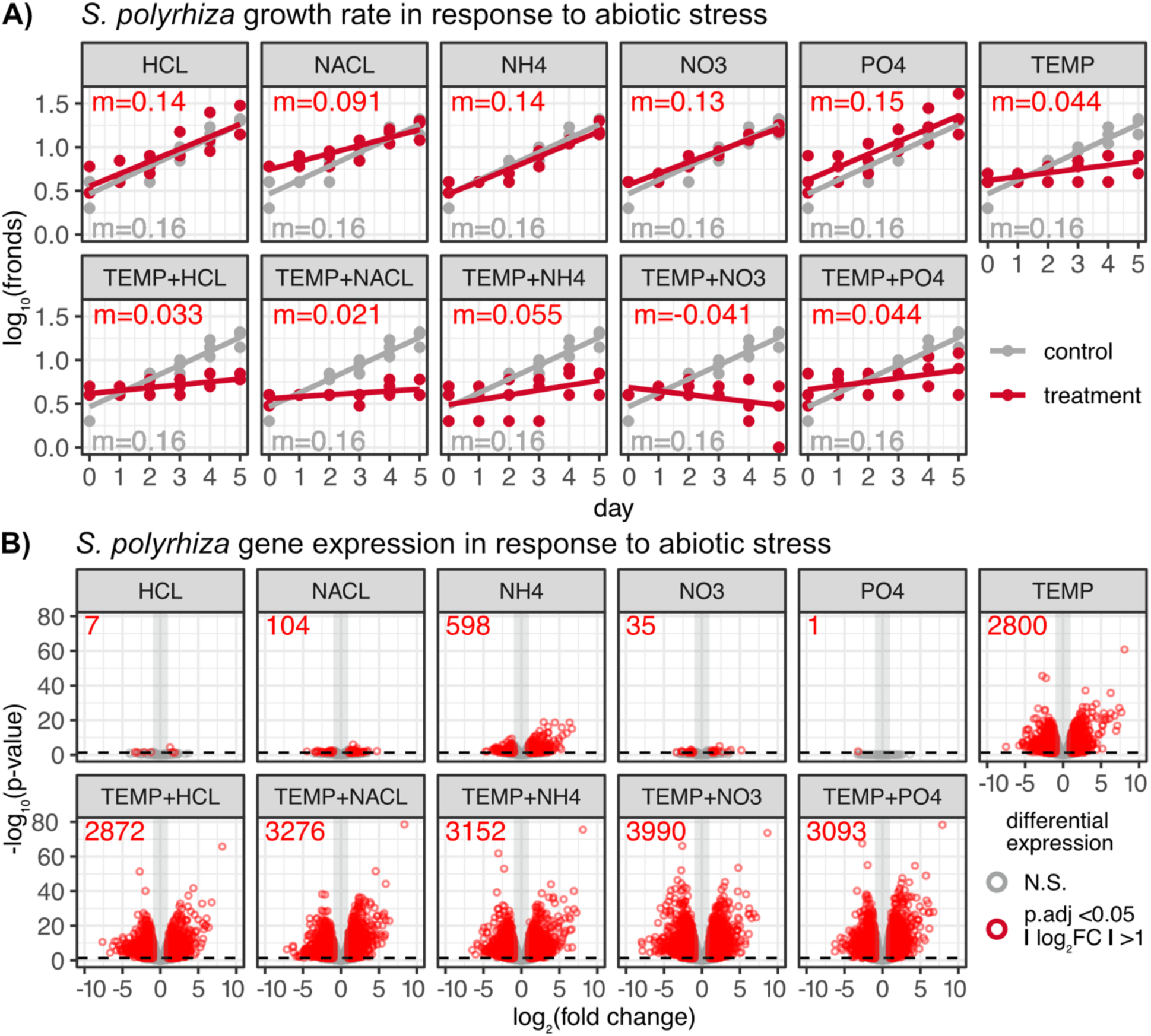
The effects of abiotic stress on the growth rate and the transcriptome of *S. polyrhiza*. A) Log-transformed *S. polyrhiza* fronds (y-axis) over a five-day (x-axis) individual stress treatment or combinatorial stress treatment (red). Untreated control is in gray. Slopes are indicated. B) Number of differentially expressed genes (red) in individual stress treatments (top row) and combinatorial treatments (bottom row). Genes with an adjusted p-value < 0.05 and a log2 fold-change greater than 1 were considered as differentially expressed (red circles).

Next, we generated corresponding transcriptomes for *S. polyrhiza*. Tissue was collected 3 hours post-treatment to capture the onset of transcriptional responses. For the combinatorial treatments, fronds were grown at 40°C for at least 24 hours before additional treatments were administered to minimize heat shock response signals (Materials and Methods, Supplemental Figure 3). To identify differentially expressed genes (DEGs), we aligned RNA-seq reads to the *S. polyrhiza* 9509 V1 genome and applied edgeR (lemna.org, Materials and Methods). As expected, the individual mild media treatments showed far fewer DEGs than the individual temperature treatment (Figure 2B). Overall, the combinatorial stress treatments showed a greater number of DEGs than expected from individual treatments (proportions test, Bonferroni corrected, Supplemental Table 2). Only the combinatorial treatment of high temperature and low pH (TEMP+HCL) failed to show this result.

To visualize gene expression differences among the samples, we performed multi-dimensional scaling of pairwise Euclidean distances of the top 500 most variable genes. We found that the temperature treatment showed the greatest effect on gene expression variation while the mild ammonium treatment had the second strongest effect, accounting for 42% and 14% of the total variation respectively (Figure 3A). We observed unique transcriptional profiles in response to the combinatorial stress treatments – on average, 32% of DEGs were not shared with the respective individual treatments (Figure 3B). Overall, we found that 2,245 different genes were differentially expressed only in combinatorial stress treatments (Figure 3C). Of these genes, 988 DEGs were unique to one specific combinatorial treatment. The TEMP+NO3 combinatorial treatment showed the greatest number of unique DEGs (418) despite the few DEGS (35) in the individual nitrate treatment and its minor effect on growth rate. 107 DEGs were shared among the five combinatorial treatments. Moreover, while the fraction of genes that were upregulated or downregulated was similar among DEGs in combinatorial stress treatments, DEGs shared among the combinatorial treatments tended to be downregulated (Supplemental Figure 4A-B). In contrast, genes that were unique to a specific combinatorial stress treatment did not show a directionality bias.

**Figure 3:**
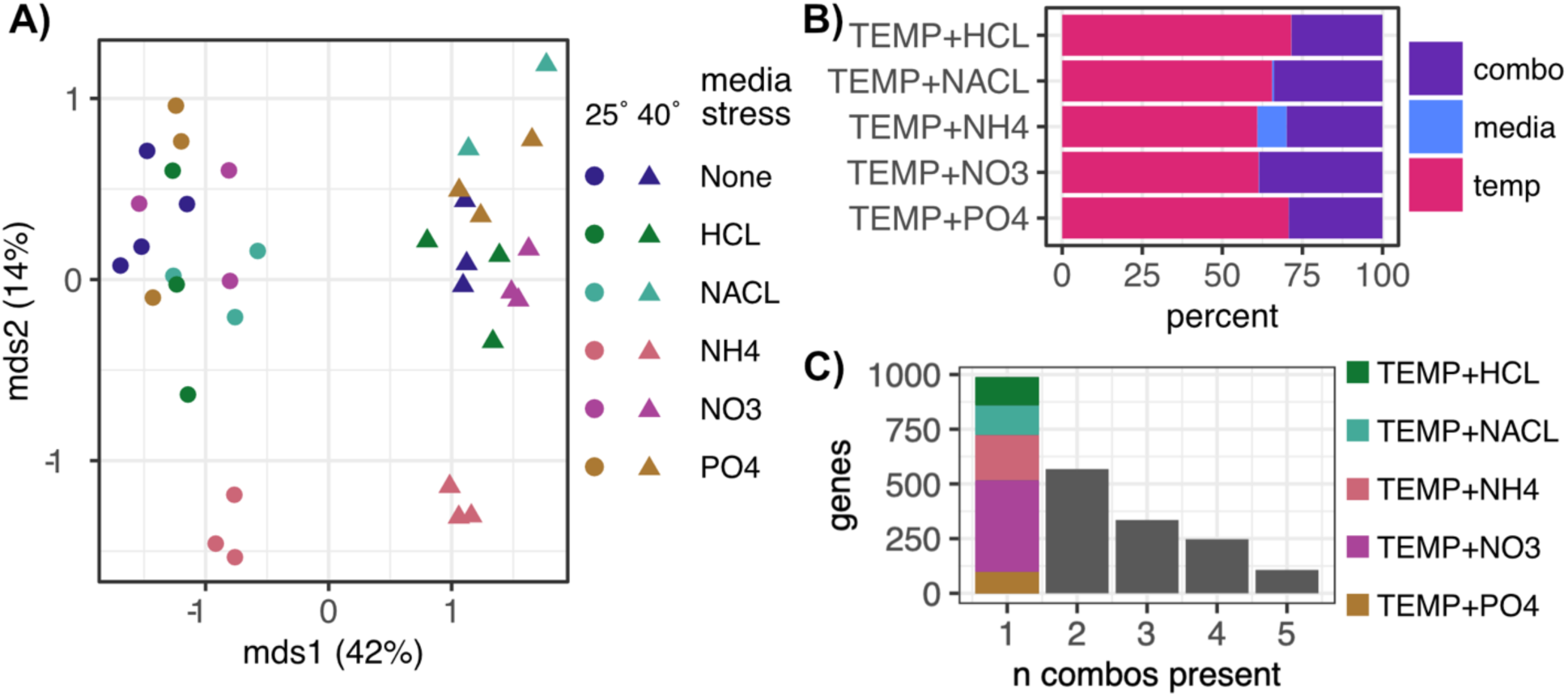
Transcriptional responses to combinatorial stress treatments are unique in *S. polyrhiza.* A) Multi-dimensional scaling using Euclidean distance of the top 500 most variable genes. The high temperature treatment explains 42% of the gene expression variation among samples (mds1) while 14% (mds2) is explained by the ammonium treatment. B) Proportions of DEGs in combinatorial stress treatments that are shared with individual treatments or unique to a combination. DEGs shared with the temperature treatment average at 66% (magenta), DEGs shared with media stress average at 2% (blue) and DEGs unique to the combinatorial treatments average at 32% (purple). C) The number of combination-specific DEGs (y-axis, total 2,245 genes) stratified by the number of combinations in which they are present (x-axis). Overall, 988 DEGs were unique to one combination; 568 were shared by two combinations; 335 by three combinations; 247 by four combinations and 107 DEGs were shared by all five tested combinatorial treatments. Of the genes unique to one combination, the combinatorial treatment with nitrate showed 418 unique DEGs. Ammonium, salt, pH and phosphate combinations showed 206, 135, 130 and 99 unique DEGs respectively.

Recent studies in *A. thaliana* identified at least 13 transcription factors involved in the response to combinations of environmental signals; certain F-box proteins have also been implicated in stress response networks (Gonzalez *et al*. 2017, Zandalinas *et al*. 2021, Sinha *et al*. 2025, Johnson *et al*. 2026, Supplemental Figure 4C). We used published data to assign *S. polyrhiza* genes to homologous orthogroups with the 13 *A. thaliana* transcription factors (TFs) (Supplemental Figure 4D). Ten *S. polyrhiza* genes in homologous orthogroups with seven of the 13 *A. thaliana* TFs showed altered expression (NAC46: AT3G04060, MYB43: AT5G16600, RAP2.3: AT3G16770, bHLH35: AT5G57150, LBD37: AT5G67420, PLATZ1: AT1G21000, bHLH-like: AT1G10585, Supplemental Figure 4E). Four *S. polyrhiza* genes in homologous orthogroups with the *A. thaliana* TFs RAP2.3 and PLATZ1 were upregulated across multiple conditions. Expression of the *S. polyrhiza* genes corresponding to the *A. thaliana* TF MYB43 (Sp9509d007g009480) was specific to the ammonium response with increased expression in the individual treatment (NH4) and the combinatorial treatment with high temperature (TEMP+NH4); the gene was downregulated in the other combinatorial treatments.

We performed GO term enrichment analysis of DEGs shared across all combinations (107 DEGs) and DEGs specific to a given combinatorial treatment (TEMP+NO3: 418, TEMP+NH4: 206, TEMP+NACL: 135, TEMP+HCL: 130 and TEMP+PO4: 99 DEGs, Materials and Methods). Across biological processes, molecular functions and cellular components, shared DEGs were enriched for terms involved in photosynthesis and the mitotic cell cycle (Supplemental Figure 5, 6 and 7). As one might expect, for DEGs present in only one combinatorial treatment, enriched terms were largely non overlapping.

### Determining the regulatory landscapes of the three duckweed species

Next, we assayed the regulatory landscape of *S. polyrhiza* with ATAC-seq (Assay for Transposase-Accessible Chromatin) in control conditions (Buenrostro *et al*. 2015, Materials and Methods). Because studies in *A. thaliana* and *Z. mays* show that chromatin accessibility is largely static across environmental conditions, developmental stages and genetic backgrounds (Sullivan *et al*. 2014, Alexandre *et al*. 2018, Sullivan *et al*. 2019, Dorrity *et al*. 2021, Paterson and Queitsch 2024, Bubb *et al*. 2025), we refrained from assessing chromatin accessibility across the tested abiotic treatments. Specifically, we wanted to examine whether the *S. polyrhiza* regulatory elements associated with differentially expressed genes shared features found for elements near environmentally responsive genes in *A. thaliana*. We also collected ATAC-seq data from *L. minor* and *W. australiana* species to distinguish between regulatory landscape features specific to *S. polyrhiza* and those specific to duckweeds.

In the three tested species, an average of 83% (SD=7.8%) of reads mapped to the respective nuclear genome and coverage was consistent (Supplemental Figure 8 and 9A). We used a heuristic peak caller to identify accessible chromatin regions in each sample and calculated the fraction of reads within peaks (FriP), an established measure of data quality (Materials and Methods). FriP scores averaged at 46% (SD=3%) for *S. polyrhiza*, 33% (SD=2%) for *L. minor* and 43% (SD=0.7%) for *W. australiana* (Bubb and Deal 2020, Material and Methods). We also determined the fraction of reads in transcription start sites (FriT) score, a more stringent quality measure, for all samples. FriT scores averaged at 25% (SD=3%) for *S. polyrhiza*, 21% (SD=1%) for *L. minor* and 18% (SD<1%) for *W. australiana*. Replicate correlation for samples was greater than r=0.88 for *S. polyrhiza*, r=0.98 for *L. minor* and r=0.95 for *W. australiana* (Supplemental Figure 9B).

Next, we determined the extent to which chromatin accessibility near the transcriptional start site (TSS) correlated with gene expression (Alexandre *et al*. 2018, Bubb *et al*. 2025). To do so, we stratified genes by expression deciles and quantified insertions per base pair in the 400 bp upstream of the TSS. The top 10% of expressed genes showed on average three-fold higher accessibility than the bottom 10%, with intermediate expression deciles showing intermediate levels of accessibility, respectively (Supplemental Figure 9C). This result, taken together with the quality metrics discussed, is consistent with high-quality chromatin accessibility data for all three duckweed species (Bubb and Deal 2020).

### Helixer and MAKER annotations yield consistent results

A common obstacle when working with emerging or non-model organisms is the lack of well-annotated reference genomes that allow accurate comparisons among species. One way to overcome this obstacle is the generation of *ab initio* annotations. We used Helixer, a framework using Deep Neural Networks and a Hidden Markov Model (Stiehler *et al*. 2020), to generate *ab initio* gene predictions for the three duckweed species. We then compared the Helixer gene predictions with those published by Ernst *et al. 2025*, which were generated with MAKER, a genome annotation pipeline using expressed sequence tags (ESTs, Cantarel *et al*. 2008, Supplemental Figure 10A and B).

We aligned our RNA-seq reads to the *S. polyrhiza* genome using both Helixer and MAKER annotations and found no significant differences in the mappability of reads (Supplemental Figure 10C). We also examined the distribution of read counts for each gene between annotations, and while the average counts per gene were slightly higher for the Helixer-annotated genes (Helixer mean = 60.1cpm, MAKER mean = 59.8cpm), the difference was not significant (Supplemental Figure 10D). Average chromatin accessibility in the 400 bp TSS-proximal region was consistent with both annotations, but there was a subtle difference in the peak of accessibility (Supplemental Figure 10E). In previously assayed plant species, this peak tends to reside ∼150 bp upstream of the TSS (Alexandre *et al*. 2018, Bubb *et al*. 2025). For MAKER annotations, the peak of accessibility also resided ∼150 bp upstream; however, for Helixer annotations, the peak was shifted to ∼50 bp upstream. This shift likely reflects that Helixer does not annotate multiple isoforms and TSSs, whereas MAKER allows for multiple TSSs to be called. Nonetheless, our comparisons suggest that when EST-based annotations are not available, Helixer is sufficient to gain biological insights.

### Duckweed ACR distribution is similar to those in terrestrial plants

Chromatin accessibility has been extensively surveyed in *A. thaliana, Z. mays,* and other terrestrial plants (Sullivan *et al*. 2014, Rodgers-Melnick *et al*. 2016, Maher *et al*. 2018, Alexandre *et al*. 2018, Lu *et al*. 2019, Sullivan *et al*. 2019, Bubb *et al*. 2025). Thus far, only a single data set is available for *S. polyrhiza* 7498 (Lu *et al*. 2019). To compare the regulatory landscapes of the three duckweed species assayed here, we determined a union set of accessible chromatin regions (ACRs) for each species, using the respective replicate data. For comparing these union ACRs with those of terrestrial plants, we generated union ACRs for previously published ATAC-seq data from *A. thaliana* and *Z. mays* (Alexandre *et al*. 2018, Bubb *et al*. 2025). The number of union ACRs called across species was similar: 27,466 ACRs in *S. polyrhiza*, 29,124 in *L. minor*, 33,823 in *W. australiana,* 34,661 in *A. thaliana* and 35,932 in *Z. mays.* The previously published *S. polyrhiza* 7498 showed 23,472 ACRs (Supplemental Table 3). Thus, despite large differences in organismal complexity and genome size, we find similar numbers of active regulatory elements in duckweed genomes compared to terrestrial plants.

Next, we quantified the total proportion of genomic features such as the TSS-proximal region (400 bp upstream of the TSS), the 5’-UTR, the 3’-UTR, exons, introns and intergenic regions for each genome and each set of ACRs across the three duckweeds, *A. thaliana* and *Z. mays* (Materials and Methods, Figure 4A-B). The examined genomes differ vastly in the total number of base pairs contributing to each genomic feature. The smallest genomes, *A. thaliana* and *S. polyrhiza*, resemble each other in the distribution of ACRs, while the larger duckweed genomes show a greater fraction of base pairs residing in intergenic regions. As expected, intergenic regions dominate the larger maize genome. Despite these stark differences in genome composition, ACRs were strongly enriched in the TSS-proximal, 5’-UTR and 3’-UTR regions across all five species and depleted in intergenic regions (Figure 4C). We observe differences in the enrichment of ACRs in coding and intronic regions – the species with small genomes show ACR depletion in these features, presumably as a direct consequence of genome size rather than functional differences.

**Figure 4.**
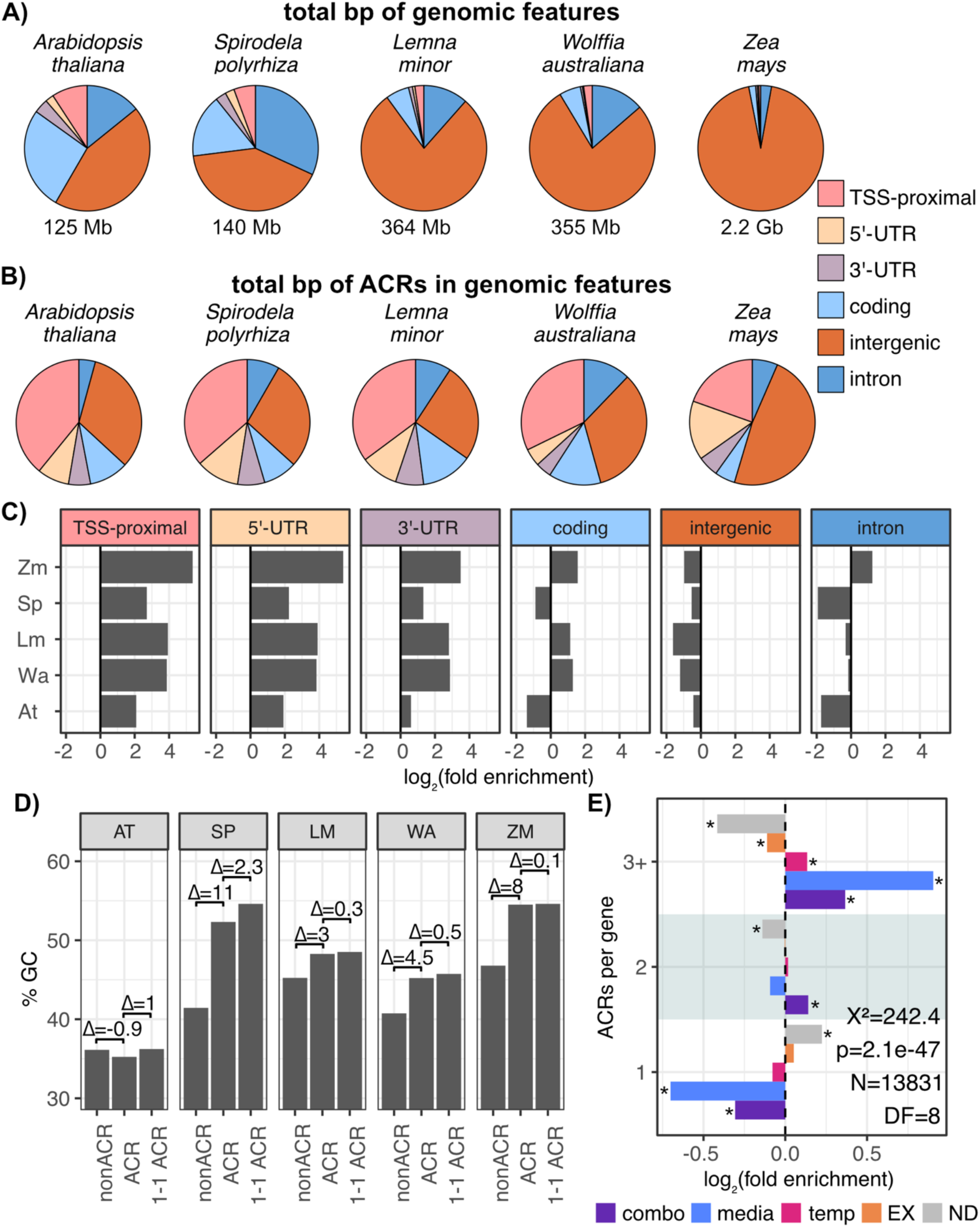
The regulatory landscapes of duckweeds mirror those of terrestrial plants. A) Total number of base pairs corresponding to genomic features in the genomes of three duckweed species, *Arabidopsis thaliana* and *Zea mays*. B) Total number of base pairs in ACRs that overlap with genomic features in the three duckweed species, *Arabidopsis thaliana* (Alexandre *et al*. 2018) and *Zea mays* (Bubb *et al*. 2025). C) The log_2_ fold-enrichment (x-axis) of the total number of base pairs of each feature in ACRs for *A. thaliana* (At), *S. polyrhiza* (Sp), *L. minor* (Lm), *W. australiana* (Wa), and *Z. mays* (Zm). Enrichments show the prevalence of TSS-proximal active regulatory elements across plant genomes of vastly different sizes, in addition to implicating genome size in differential enrichments. D) GC content of the five plant species in non-accessible regions (nonACR), accessible chromatin regions (ACR), and ACRs proximal to genes with one-to-one orthologs (1-1 ACR). *S. polyrhiza* showed the greatest increase in GC content in ACRs compared to nonACR GC content. E) The log_2_ enrichment of *S. polyrhiza* genes in five categories (differentially expressed in high temperature (temp), differentially expressed in media treatments (media), differentially expressed in a combinatorial treatment (combo), expressed genes (EX) and genes that were not detected (ND). Genes are stratified by the number of associated ACRs (1, 2 or 3+). Chi-square test of independence (X^2^=242.4, p=2.1e-47, N=13831, DF=8). Asterisks indicate features with significant standardized residuals from Chi-square test (| std residual | > 2.0, p < 0.05).

Further, we compared GC content in genomes and ACRs across species (Figure 4D). As expected, genome GC content was higher in all monocot species compared to the dicot *A. thaliana*. GC content in monocot genomes was even higher in ACRs with *S. polyrhiza* showing the greatest difference in %GC (Δ=11%) between genomic non-ACR regions and ACRs, even greater than the difference observed in *Z. mays* (Δ=8%). *A. thaliana* ACRs showed the known decrease in GC content compared to the genome itself (Jores *et al*. 2021, Jores *et al*. 2026). These trends held true when examining ACRs near genes that are 1 to 1 orthologs. The high GC content of regulatory regions in maize has sometimes been speculated to be a protection mechanism against the insertion of transposable elements. Our results challenge this theory as the *S. polyrhiza* genome contains the fewest transposable elements among the examined duckweeds and does not differ substantially in transposon content from the dicot *A. thaliana*.

### Environmentally responsive genes are enriched for being associated with multiple ACRs

Previous studies in *A. thaliana* show that environmentally responsive genes tend to be associated with multiple ACRs (Sullivan *et al*. 2019). To verify if this association holds true in duckweed, we classified *S. polyrhiza* genes into one of five categories: (1) differentially expressed in high temperature; (2) differentially expressed in a media treatment; (3) differentially expressed in combinatorial conditions; (4) detected but not differentially expressed or (5) not detected in our expression data. Genes in categories 1-3 were considered environmentally responsive genes. Of the 20,342 annotated genes, 2,800 genes fell into the first category, 399 into the second, 2,243 into the third, 9,795 into the fourth and 5,103 into the last. We assigned all genes into categories based on their number of associated ACRs (1, 2 or 3+ associated ACRs (Materials and Methods, Supplemental Figure 11A) and calculated log_2_ fold enrichment of genes in expression and ACR categories. Consistent with the prior *A. thaliana* findings, we find the environmentally responsive genes are enriched for being associated with 3+ ACRs and depleted for being associated with a single ACR (Figure 4E). Among environmentally responsive genes, media-responsive genes (category 2) showed the greatest enrichment for 3+ ACRs. Category 1 genes showed a modest but significant enrichment for 3+ ACRs but no depletion for single ACRs. In contrast, genes that were not differentially expressed in any of the conditions (category 4) were depleted for being associated with 3+ ACRs, while being enriched for association with single ACRs.

Next, we asked if this pattern holds for orthologous genes in the *L. minor* and *W. australiana* genomes, grouping categories 1-3 together as differentially expressed genes (Materials and Methods, Supplemental Figure 11B, Earnst *et al*. 2025). We inferred 4,861 genes in *L. minor* and 4,161 genes in *W. australiana* as differentially expressed, 6,975 and 6,524 genes as expressed, and 1,227 and 1,006 genes as not detected, respectively, and calculated log_2_ enrichment of genes with 1, 2 or 3+ associated ACRS. *L. minor* and *W. australiana* genes inferred to be environmentally responsive were enriched for being associated with 3+ ACRs and depleted for single ACRs (Supplemental Figure 11C). Genes inferred to be not responsive or not expressed were depleted for multiple ACRs and enriched for single ACRs. We conclude that environmentally responsive genes in duckweeds tend to be associated with multiple ACRs, mirroring a trend first observed in the terrestrial crucifer model *A. thaliana* (Sullivan *et al*. 2014, Sullivan *et al*. 2019).

### ACRs associated with genes differentially expressed in combinatorial stress show unique enrichments of TF binding motifs

We performed motif enrichment analysis of ACRs to identify candidate TFs driving gene expression in response to the tested treatments (Supplemental Figure 12A-B). To do so, we stratified ACRs by the direction of expression change of associated genes and used TF binding consensus motifs to examine their enrichments (Materials and Methods, McLeay & Bailey 2010, O’Malley *et al*. 2016, Jorres *et al*. 2021). Seven TF binding motifs were enriched across the examined ACRs (Figure 5A). ACRs associated with genes upregulated in individual and combinatorial treatments were enriched for the MYB/ CAMTA/ FAR1 family consensus motif. The canonical heat shock element was enriched in ACRs associated with upregulated genes in both the high temperature and combinatorial treatments despite the loss of several HSF genes in this family (Supplemental Figure 12C). The C2H2 family consensus motif was specifically enriched in ACRs associated with genes upregulated in combinatorial treatments, suggesting that TFs binding this motif might specifically act in response to multiple environmental inputs. However, orthogroup analysis did not yield evidence for *A. thaliana*-like C2H2 transcription factors in the three duckweeds (Figure 5B).

**Figure 5:**
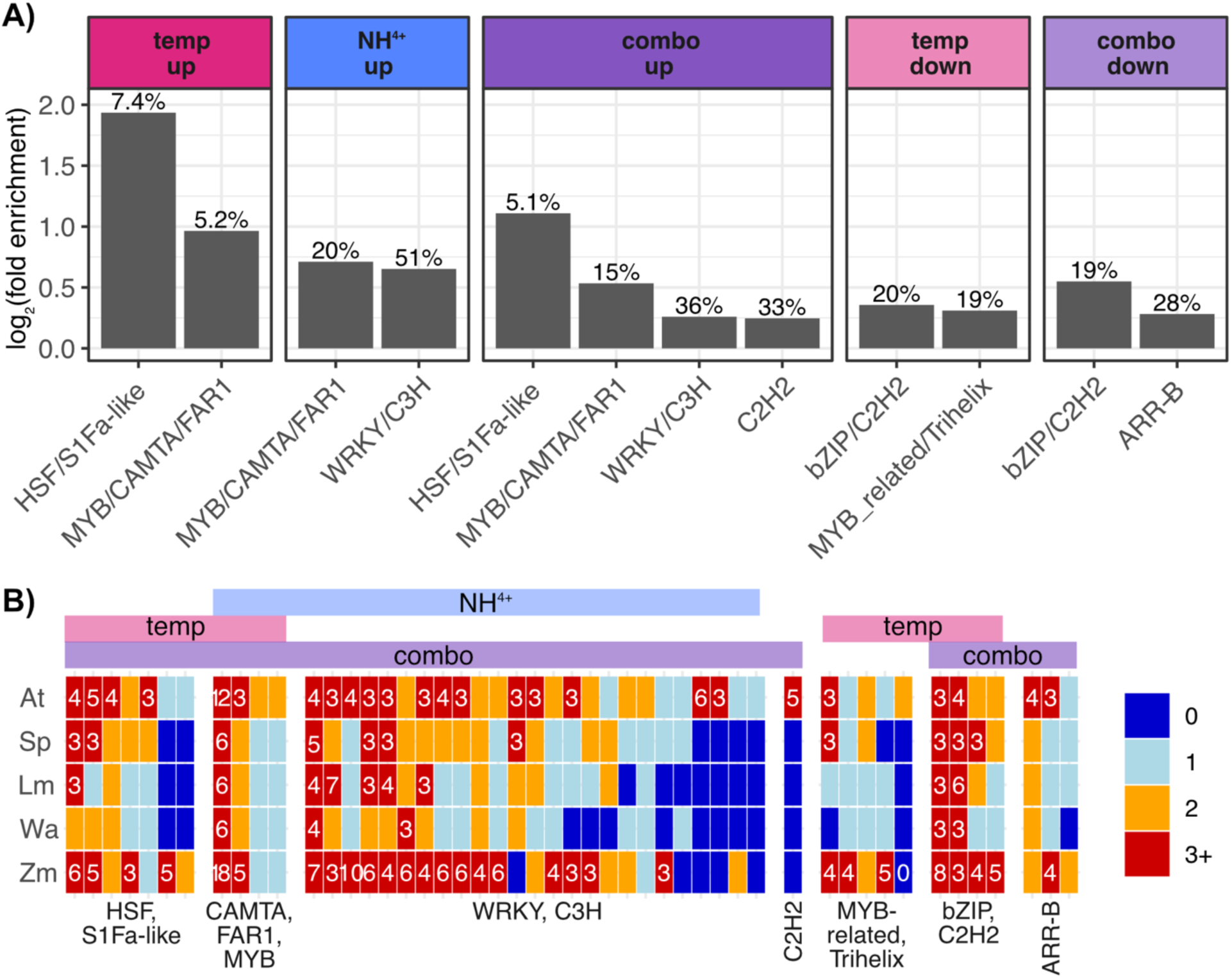
C2H2 family motif is enriched in ACRs associated with genes responsive to combinatorial treatments but orthologs are missing in duckweed. A) Log_2_-fold enrichment (y-axis) of consensus motif families (x-axis) in subsets of ACRs. Percentages of total ACRs with positive hits are displayed on top of each bar. B) Orthogroups (columns) of *A. thaliana* transcription factors genes recognizing enriched consensus motifs for the examined species (rows). Orthogroups are grouped by motif family (bottom labels); the motif families are ordered by the subset of ACRs in which they are enriched in (top labels).

## Discussion

In plants, the study of combinatorial stress treatments has been limited by the size and generation time of existing models. The small and rapidly dividing duckweeds are well suited for high throughput and multiplexed studies and offer the opportunity to study plant stress on a scale and combinatorial complexity not easily achievable in terrestrial plants. Here, we have examined growth responses, transcriptomes and accessible chromatin regions in duckweeds to evaluate their potential to serve as models for terrestrial plants.

Prior studies found that combinations of abiotic stress treatments result in unique expression responses in terrestrial plants that are not predictable from the responses to the individual treatments (Ristova et al. 2016, Zandalinas et al. 2021). We find the same to be true for the duckweed *S. polyrhiza.* The transcriptomes measured across five combinatorial treatments showed that on average 32% of DEGs were not shared with the corresponding individual treatments. This result is remarkable considering that the combinatorial treatments all included increased temperature, which yielded by far the strongest transcriptional response as an individual treatment. Further, almost 1000 DEGs were unique to one of the tested combinations. We conclude that the *S. polyrhiza* response to combinatorial abiotic stress resembles those found in terrestrial plants: first, response to individual treatments does not predict the response to combinatorial treatments, and second, combinatorial responses are largely unique.

To date, the regulatory landscape of duckweeds has only been assayed in a single species (*S. polyrhiza* accession 7498, Lu *et al*. 2019). Here, we add chromatin accessibility data for a second strain of *S. polyrhiza* (accession 9509), for *W. australiana* (accession 8730), and for *L. minor* (accession 7210). We observed similar numbers of ACRs in the duckweeds to those found in the small *A. thaliana* genome and in the vastly larger maize genome. Thus, somewhat surprisingly, the number of ACRs did not appear to reflect genome size or organismal complexity, at least when assayed in whole duckweed fronds and compared to ACRs found in *A. thaliana* seedlings and maize leaves.

However, genome size was subtly reflected in the enrichments of ACRs in genomic compartments. In all species, regardless of genome size, the largest enrichment of ACRs was found within 400 bp upstream of TSSs while ACRs were depleted in intergenic regions. The species with the smallest genomes, *A. thaliana* and *S. polyrhiza,* were also depleted for ACRs in coding and intronic regions and closely resembled each other. Similarly, the ACR enrichments of *W. australiana* and *L. minor*, the two duckweed species of nearly identical intermediate genome size, resembled one another. *Z. mays* was the only species to show ACR enrichment in introns.

Elevated GC content is a major difference between monocot and dicot genomes; this difference appears to be amplified in ACRs in the species examined to date (Jores *et al*. 2021, He *et al*. 2024, Jores *et al*. 2026). *S. polyrhiza* shows the highest GC content of the three examined duckweed species, comparable to that of *Z. mays*, and exhibits the largest difference in GC content between ACRs and the genome of all the examined species. High GC content in the regulatory regions of monocots is sometimes invoked as a protective mechanism against the insertion of transposable elements that preferably target ACRs and regions with diffuse chromatin accessibility (Liu *et al*. 2009, Stitzer *et al*. 2021, Boissinot 2022, Bubb *et al*. 2025, Scherer & Nelms 2026). *S. polyrhiza’s* transposon load is comparable to that of *A. thaliana* (Wang *et al*. 2014, Dombey *et al*. 2025), seemingly refuting this hypothesis. However, the reduction of RdDM-associated genes in duckweeds might render their transposable elements more active (Dombey *et al*. 2025, Ernst *et al*. 2025).

Taken together, our data provides evidence that *S. polyrhiza* shares fundamental features with terrestrial plants, including monocots with large genomes like maize. We found strong similarities in the response to combinatorial stress treatments, in the distribution of ACRs across genomic compartments, in the association of multiple ACRs with environmentally responsive genes and in ACR GC content with terrestrial counterparts. Hence, these small monocots with their highly simplified body plans are appropriate models for disentangling the complex mechanisms underlying combinatorial stress responses in the complex monocots that contribute to our food supply.

## Materials and Methods

### Duckweed stocks and growth conditions

Duckweed lines were obtained from the Rutgers Duckweed Stock Cooperative (www.ruduckweed.org). For this study *Spirodela polyrhiza* 9505, *Lemna minor* 7210, and *Wolffia australiana* 8730 were maintained at 25°C 16 h light/8 h dark; 300 µmol m^-2^ sec^-1^, 25°C, 70% relative humidity in Conviron Chambers. Fronds were grown in liquid 0.5 X Schnek and Halbred media with 0.05% sucrose and 100µg/L cefotaxime in 100x 25mm deep petri dishes.

### Growth assays

Growth assays were performed by transferring fronds into 6-well plates with 7ml of test media. Photos were taken every day at ZT-8 for five days and the total number of live fronds were counted. To calculate growth rates (slope), frond counts were log10 transformed and fit to a linear model (Supplemental Code, Supplemental Data 1-2) and the slope of the best fit line was extracted. Samples without living fronds were removed. To determine the range of stress tolerance to individual treatments, duckweed were challenged with different intensities of six individual stressors: ammonium overabundance (NH4: ammonium acetate 1M, 0.2M, 0.02M), salt overabundance (NACL: sodium chloride 1M, 0.1M, 0.05M), phosphate overabundance (PO4: potassium phosphate 1M, 0.2M, 0.02M), nitrate overabundance (NO3: potassium nitrate 1M, 0.1M, 0.05M), acidity (HCL: 2,3,4,5), or heat stress (TEMP: C 30°, 35°, 40°). Combinatorial testing was done using two stress conditions at time, one mild media stress (NH4: 0.02M, NaCl: 0.05M, PO4: 0.02M, NO3: 0.05M or HCL: 4) and a temperature stress (TEMP: 40°C).

### Transcriptome sequencing

For transcriptome sequencing, we sampled individual stresses (NH4: 0.02M, NACL: 0.05M, PO4: 0.02M, NO3: 0.05M, PH: 4 and TEMP: 40°C) as well as combinatorial stresses consisting of one media stress and one temperature stress in *S. polyrhiza*. For treatment conditions involving high temperature, fronds were first acclimated to high temperature for 24 hours to minimize signals from the heat shock response. To capture the onset of transcriptional response, samples were collected 3 hours after exposure to challenge conditions and snap frozen in liquid nitrogen before extraction. RNA was extracted from samples in seven batches of random subsamples to prevent confounding signal from the RNA extraction, using the QIAGEN RNeasy Plant Mini Kit (Cat: 74904). Sequencing libraries were prepared using Illumina stranded mRNA Prep Ligation kit (Cat: 20040532). We then performed paired end library sequencing using the Illumina NextSeq 2000 platform, with 50bp paired end reads.

### Genome annotations

Genome and genome annotations for *Spirodela polyrhiza* 9505 V2, *Lemna minor* 7210 V1, and *Wolffia australiana* 8730 V2 were downloaded from lemna.org (Ernst *et al*. 2025). However, the *W. australiana* V2 assembly did not have a corresponding annotation, and so the *W. australiana* V1 annotation coordinates were lifted over to the V2 genome using the UCSC Liftover tool (Hinrichs *et al*. 2006). A*b initio* genome annotations for all three duckweed genomes were generated with the Helixer online utility (Steihler *et al*. 2020).

Sequenced RNA libraries were aligned to the *Spirodela polyrhiza* V2 genome using STAR read aligner (Dobin *et al*. 2013, 2.7.11b, run parameters *--runThreadN 10 –-quantMode TranscriptomeSAM GeneCounts –-outSAMstrandField intronMotif –-twopassMode Basic –-readFilesCommand zcat*). For the 37 generated libraries across 12 treatment conditions and three biological replicates, sample meta data, MAKER and Helixer count tables were downloaded into R (R Core Team 2025, Posit team 2024, 4.4.2, Supplemental Data 3-5). Only one library was excluded due to low read counts, this library corresponded with the TEMP+NACL combinatorial challenge, leaving this condition with two replicates. Differential gene expression analysis was performed using edgeR (4.4.2), extraction batches were included in the design matrix and visualization was performed using ggplot2 (3.5.2, Chen *et al*. 2025, Wickham 2016, Supplemental Code).

### Gene Ontology (GO) analysis

For GO analyses, we first generated a functional annotation using the Blast2GO (Götz *et al*. 2008) functionality within OmicsBox (3.5). GO term enrichment analysis was then performed using R package topGO (Alexa & Rahnenführer 2025, 2.58.0, Supplemental Code). P-values from Fischer’s exact test of independent GO terms, were Bonferonni corrected and significant terms plotted using ggplot2 (*runTes*t parameters: *algorithm=“classic”, statistic=“fisher”*).

### Chromatin accessibility sequencing

To quantify the chromatin accessibility of the three duckweed species, nuclei were extracted from fresh duckweed samples grown in control conditions. Fronds were first finely chopped in weigh boats on ice with 2ml of grinding buffer (0.8M sucrose, 10mM MgCl, 25mM Tris-HCl pH 8.0) and 1x protease inhibitor (cOmplete Protease Inhibitor Cocktail – Millipore Sigma: 04693132001). Samples were then transferred into gentleMACS C tubes and ground using the gentleMACS dissociator system (program e1.1 and e1.2). Extract was filtered through miracloth and centrifuged at 2000g for 5 min 4°C and the resulting pellet was resuspended in 750 µl of suspension buffer (0.4M sucrose, 10mM MgCl2, 25mM Tris-HCl pH 8.0, 0.1% Triton X). Suspension was then layered over 750 µl of 75% Percol buffer (0.4M sucrose, 10mM MgCl2, 25mM Tris-HCl pH 8.0, 75% Percol by volume, 1x protease inhibitor) and centrifuged at 2500g for 15min 4°C. Percol and interphase layers were collected and diluted in suspension buffer then spun down once more at 1500g for 5min 4°C. Final nuclei pellets were resuspended in 100µl of suspension buffer. Total nuclei per sample were calculated visually using hemocytometer and 50,000 nuclei were used per Tn5 reaction. Tagmentation reactions were set up as 50µl reactions with 25µl of 2x tagmentation buffer (20mM Tris-HCl pH7.6, 10mM MgCl, 20% vol/vol DMF), 2.5µl of 100mM loaded Transposase (Diagenode: C01070012) and 22.5µl of water. Libraries were barcoded using Nextera Index Kit primers and KAPA HiFi DNA polymerase. Sequencing of libraries was done on the Illumina NextSeq 2000 platform with 50 bp paired end sequencing.

### Read processing and quality metrics

ATAC-seq reads from each species were aligned to their respective genomes using bwa aln (BWA: 0.7.17, Li 2013) with default the parameters. For each species base-pair resolution genome coverage was calculated using bedtools *genomecov* function and then mean coverage for 1 Mb windows across the genome. Z-scores were calculated for each window by subtracting the coverage value by the mean coverage of each species then dividing by the standard deviation. Outliers were classified as having an absolute value of Z-score greater than 3. Using samtools (1.19, Danecek *et al*. 2021), reads with a quality mapping score of 60 or higher were selected to quantify total cuts per base pair position. A sliding window histogram of Tn5 insertions was generated for the entire genome (150 bp window; 20 bp slide) in the format of a bed file. 150 bp windows were labeled as peaks if they satisfy the following criteria: (1) the Tn5 insertion count within the window was in the top *p* percentile, and (2) the window had the highest Tn5 insertion count of the previous and subsequent five windows. The percentile *p* chosen scaled with genome size, although not linearly, to result in a similar number of peaks called for different-sized genomes (Bubb & Deal 2020).

Peak calls for each sample were then used to calculate the fraction of reads within peaks (FriP) for individual samples. We also calculated the fraction of reads near the transcriptional start site (FriT) scores for both MAKER and Helixer annotations, using the 400 bp upstream region of the TSS. Replicate correlations were performed by merging read files across replicates and calling peaks using the merged read file. Coordinates from the resulting peaks were then used to calculate the number of reads per peak for each replicate and the resulting count tables were correlated using Pearson correlation.

To generate expression stratified plots for *S. polyrhiza* accessibility, we used the transcriptome data and the MAKER annotations to stratify genes into expression deciles. For each decile, we then calculated the number of inserts at each position of the 400 bp upstream region of the TSS. The number of inserts at each position was normalized by the total number of inserts of the sample in order to compare across replicates.

### Analysis of ACRs

Coordinates from the union of peaks across biological replicates were used to call species-specific accessible chromatin regions (ACRs) using bedtools *merge* (2.31.1, Quinlan & Hall 2010). Called ACRs were assigned to the nearest gene feature using BEDOPS *closest-feature* with the parameters *--delim ‘\t’ –-dist –closest*. Region overlap of ACRs to genomic features was calculated using *bedmap* with parameters –-*echo –-echo-overlap-size –-delim ‘\t’* (v2.4.41, Neph *et al*. 2012). Generated tables were then imported into R for analysis and visualization using ggplot2. Enrichment of genomic features represented in ACRs was calculated as:

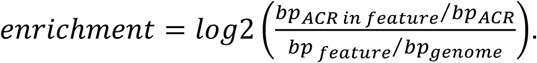

To calculate enrichment of ACRs near genes, we first assigned genes into one of five expression categories using RNA-sequencing measurements. Genes with a CPM < 1 were labeled as not detected, genes that were detected at a CPM > 1 but were not differentially expressed were categorized as expressed, differentially expressed genes were categorized as being singleton media stress induced, singleton temperature induced or combinatorially induced. For each gene we then assigned ACR categories based on the number of nearby ACRs as determined by the BEDOPS. Genes with only a single nearby ACR were categorized as “1”, genes with two associated ACRs as “2” and genes with three or more associated ACRs were categorized as “3+”. Enrichment for ACR number among expression categories was calculated as:

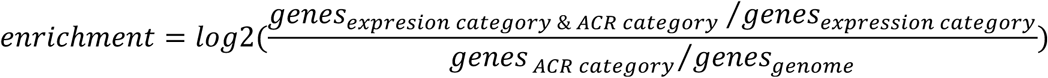

(Supplemental Figure 11A). To test for significance in enrichment a Chi-square test of independence was performed on contingency tables from using R base function *chisq.test*. Features were considered significantly different from expected distribution if the p-value of the Chi-square test was less than 0.05 and the absolute value of the standardized residual for a feature was greater than 2.

To calculate enrichment of ACRs in *L. minor* and *W. australiana*, for which we did not have corresponding transcriptome data, we used a homology-based approach to assign expression categories to genes. We took previously published homologous orthogroups (HOGs) for *S. polyrhiza*, *L. minor,* and *W. australiana* and assigned *S. polyrhiza* categories to each HOG (Earnst *et al*. 2025, Supplemental Figure 11B). For this analysis, we did not sub-divide differentially expressed genes. In cases where HOGs contained *S. polyrhiza* genes from multiple expression categories, assignment priority was given to differentially expressed genes, then expressed genes. These categories were then used to calculate enrichment of ACR number in genes from each expression category for *L. minor* and *W. australiana*.

### Motif analysis

To analyze *S. polyrhiza* ACR sequences for enrichment of motifs, we first selected *S. polyrhiza* ACRs that overlapped TSS-proximal and intergenic regions (Supplemental Figure 12A). We then grouped ACRs by their association to genes in each of the five previously described expression categories. We further divided differentially expressed categories by the direction of expression (up or down). For our media responsive set, we only used ACRs near genes differentially expressed in the individual ammonium treatment, because 80% (598 genes) of all differentially expressed genes identified from the individual media treatments were from this condition alone. We made six sets of test sequences (temp up, temp down, NH4 up, NH4 down, combo up and combo down, Supplemental Figure 12B). To find enriched motifs, we used MEME-suites Analysis of Motif Enrichment tool (AME) providing ACR sequences from expressed and not detected categories as the background sequence (5.5.9, McLeay & Bailey 2010). As our motif set, we used a previously published database of Arabidopsis motifs clustered by motif similarity into consensus motifs (Jores *et al*. 2021). Consensus motifs with an E-value > 0.05 were considered significant and enrichment was calculated as:

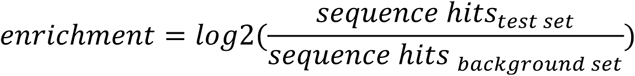

Comparison of C2H2 motif to plant (Arabidopsis; DAP motifs) vertebrate databases (Eukaryote DNA; Vertebrate) was done using MEME-suites Tomtom tool with default parameters (5.5.9, Gupta 2007).

## Author contributions/Acknowledgment

B.R-C., C.Q. & J.C. designed the research; B. R-C. & S.C. performed experiments and data collection; B.R-C. & K. B. performed the analyses, B.R-C., S.C., K.B., C.Q. & J.C. wrote the article; all authors revised and approved the article. This work was supported by the National Science Foundation (PlantSynBio Grant No. 2240888 to C.Q., Trilateral FPP No. 2519785 to C.Q., NSF Postdoctoral Research Fellowship in the Biology Program Grant No. 2305660 to B.R-C., NSF REU Award No. 2446018 to C.Q.), the National Institutes of Health (NIGMS MIRA Grant No. 1R35GM139532 to C.Q.), and the NASA BioS-ENDURES Consortium (Seed Project No. GR064401 to C.Q).

## Data Availability

Transcriptome and chromatin accessibility data can be found in the NIH short read archive (https://submit.ncbi.nlm.nih.gov/subs/sra/) under BioProject ID PRJNA1522487.

