## Supplemental Materials for "Duckweeds as Multiplexable Plant Models for Exploring Abiotic Stress Responses"

Supplemental Figures:

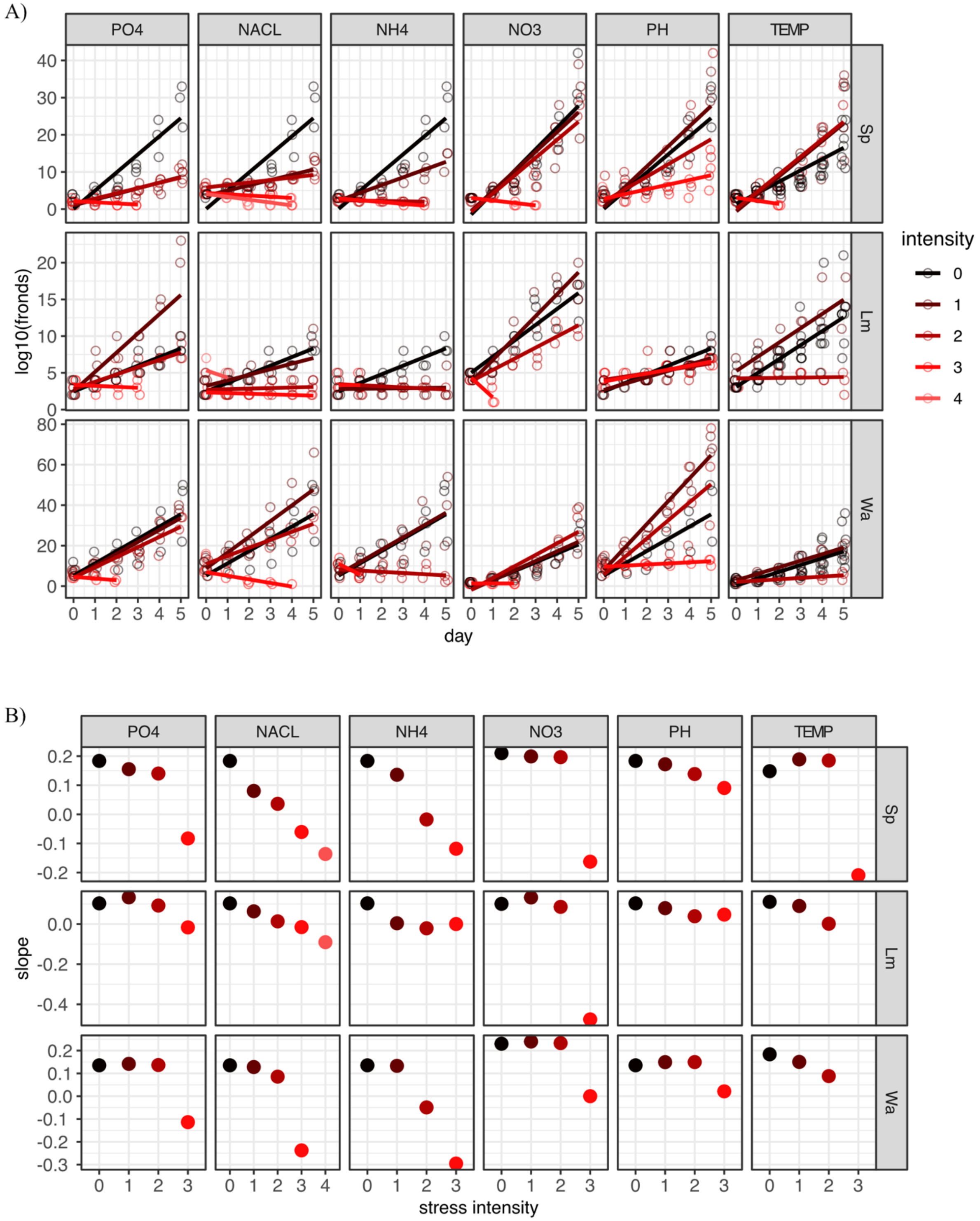


**Supplemental Figure 1: Growth rate of three duckweed species decreased with stress severity.** A) Log total fronds (y-axis) over days (x-axis) for *S. polyrhiza*, *L. minor*

and *W. australiana* in response to varying intensities of stress. Lines represent the best fit by linear model prediction. Data points without living fronds present were removed. The relative effect of individual treatments on growth was dependent on the intensity of stress and the species. B) Calculated slope (y-axis) of the best fit line for three duckweed species over increasing stress intensity (x-axis). Slopes tended to decrease with stress intensity; rates were species- and condition-specific.

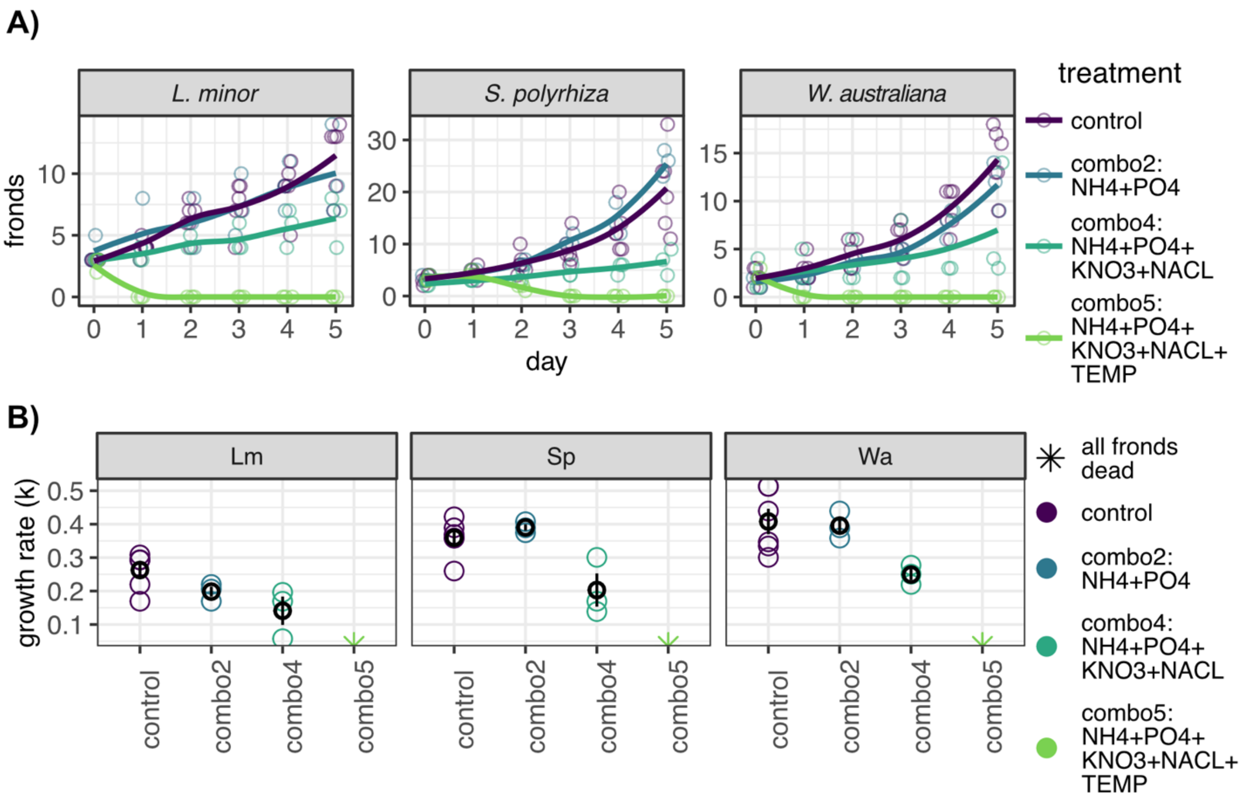


**Supplemental Figure 2: Increased stress complexity decreases growth rate in duckweeds.** A) Raw growth curves three duckweed species over a five-day time course in response to three combinatorial stresses. Total fronds on (y-axis) over a five-day time course (x-axis). B) Growth rate (y-axis) of biological replicates assayed under combinations of either two, four and five biotic stresses (x-axis). Mean growth rate and standard deviation are shown in black. Circles indicate calculated growth rate (k) asterisk indicates no living fronds which resulted in k not being computable.

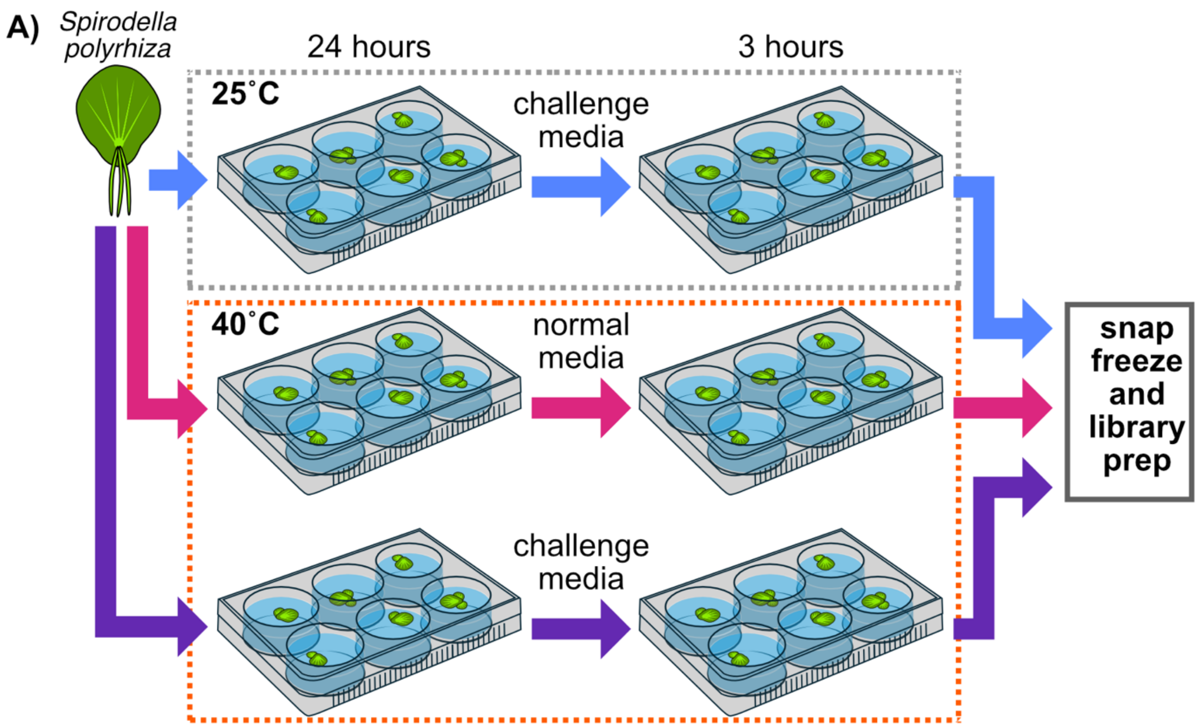


**Supplemental Figure 3: Experimental set up for individual and combinatorial treatments.** Fronds were placed into 6-well plates with normal media and placed in either 25˚C chambers or 40˚C chambers for 24 hours. Fronds were then moved to challenge media in the media treatments (blue arrow) or the combinatorial (purple arrow) treatments or moved into normal media for temperature treatment (magenta arrow).

A)

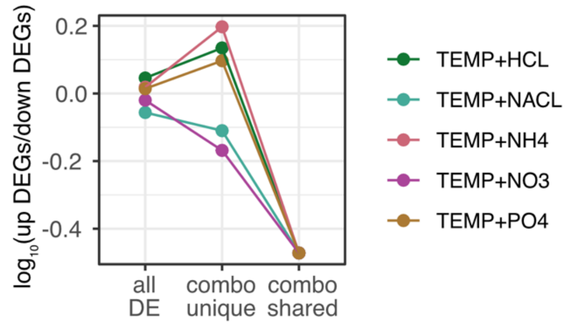


B)

| Group | combo (TEMP+) | UP | DOWN |
|---|---|---|---|
| AllDE | HCL | 1387 | 1248 |
| | NACL | 1419 | 1615 |
| | NH4 | 1450 | 1389 |
| | NO3 | 1693 | 1772 |
| | PO4 | 1467 | 1420 |
| combo unique | HCL | 75 | 55 |
| | NACL | 59 | 76 |
| | NH4 | 126 | 80 |
| | NO3 | 169 | 249 |
| | PO4 | 55 | 44 |
| combo shared | HCL/ NACL/ NH4/ NO3/ PO4 | 27 | 80 |

C)

| gene name | gene id | source |
|---|---|---|
| NLP7 | AT4G24020 | Johnson2026 |
| bHLH35 | AT5G57150 | Sinha2026 |
| bHLH-like | AT1G10585 | Sinha2026 |
| PLATZ1 | AT1G21000 | Sinha2026 |
| RAP2.6 | AT1G43160 | Sinha2026 |
| NAC46 | AT3G04060 | Sinha2026 |
| MYB-like | AT3G11280 | Sinha2026 |
| RAP2.3 | AT3G16770 | Sinha2026 |
| BBX30 | AT4G15248 | Sinha2026 |
| RD26 | AT4G27410 | Sinha2026 |
| MYB43 | AT5G16600 | Sinha2026 |
| LBD37 | AT5G67420 | Sinha2026 |
| FBOX1 | AT1G47056 | Gonzalez2017 |

D)

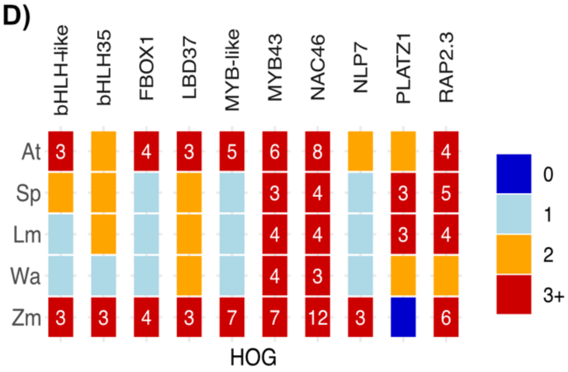


E)

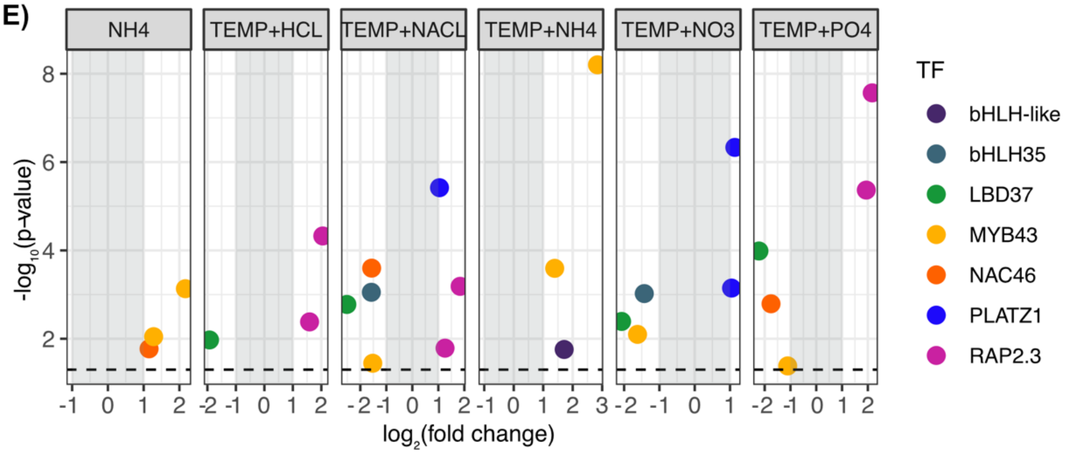

**Supplemental Figure 4: DEGs shared across combinatorial challenges were generally down regulated.** A) The log10 ratio of upregulated genes to down regulated genes (y-axis) for all differentially expressed genes (all DE), genes that are differentially expressed specifically in one combinatorial challenge (combo unique), and genes that are differentially expresses in all combinatorial treatments (combo shared). B) The number of up- and down-regulated DEGs for each group and combination C) Genes of thirteen *A. thaliana* transcription factors that were previously found to be involved in combinatorial stress responses. D) HOGs (columns) for 10 of the known genes involved in combinatorial response across 5 species (rows). Three genes (RAP2.6, BBX30 and RD26) did not have corresponding HOG data. E) Significance (y-axis) and $log_2$ fold change (x-axis) of differentially expressed genes in HOGs with TFs known to be involved in the response to combinatorial abiotic stress.

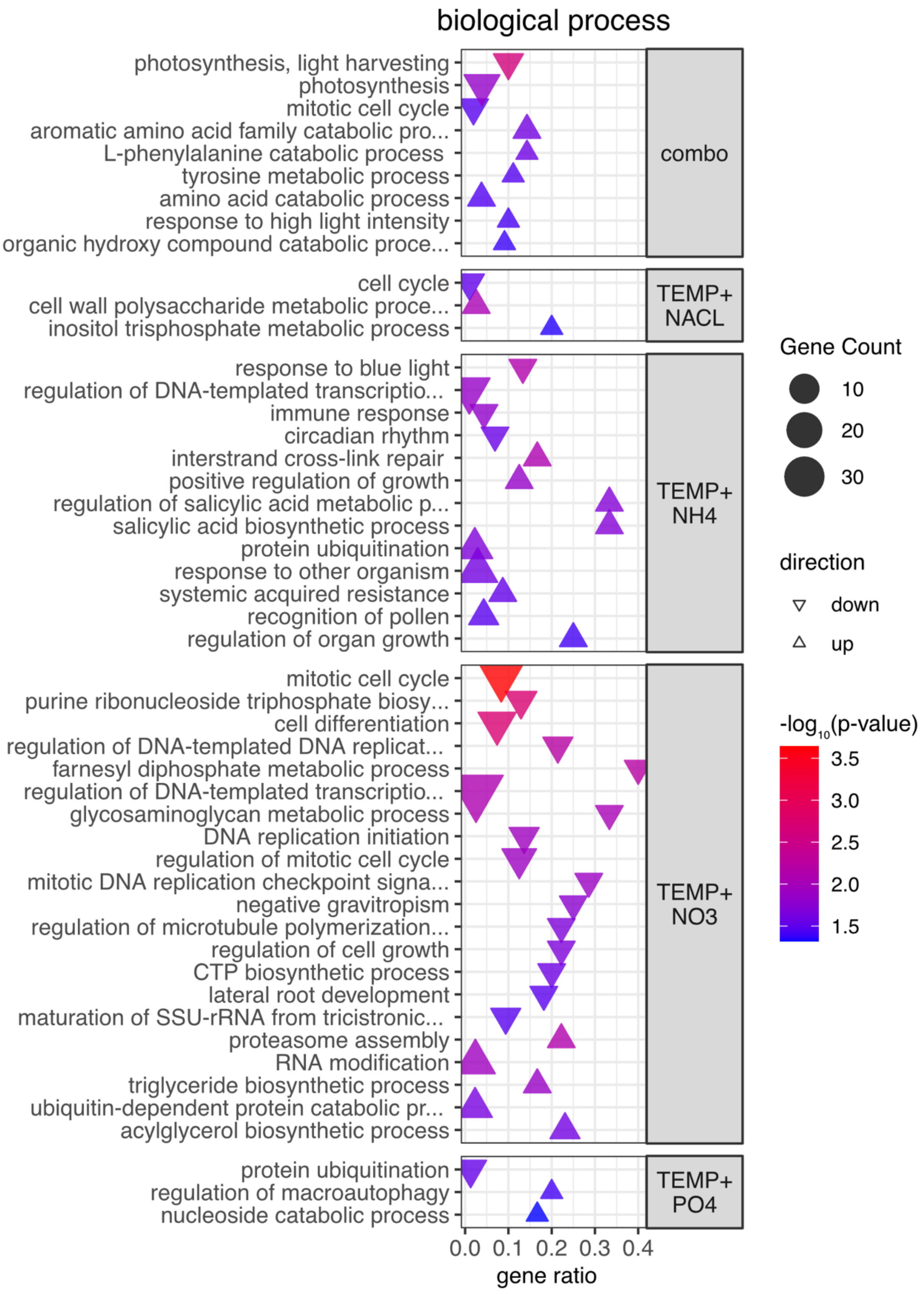


**Supplemental Figure 5: GO enrichments for biological processes suggest unique responses to combinatorial stress.** Significant GO terms for biological processes in

DEGs shared across combinatorial challenges (combo) and DEGs unique to specific combinations of stress (y-axis). Gene ratio (x-axis) indicates the proportion of the gene list enriched for a GO term. Shape size indicates total number of genes enriched for a term; upward arrow indicates GO term significance among upregulated genes while downward arrow indicates enrichment in down regulated genes. Shape color corresponds with the degree of significance after Bonferroni correction for multiple tests. GO term enrichments suggest distinct strategies are employed in response to each stress combination. TEMP+HCL treatment did not show significant GO term enrichments.

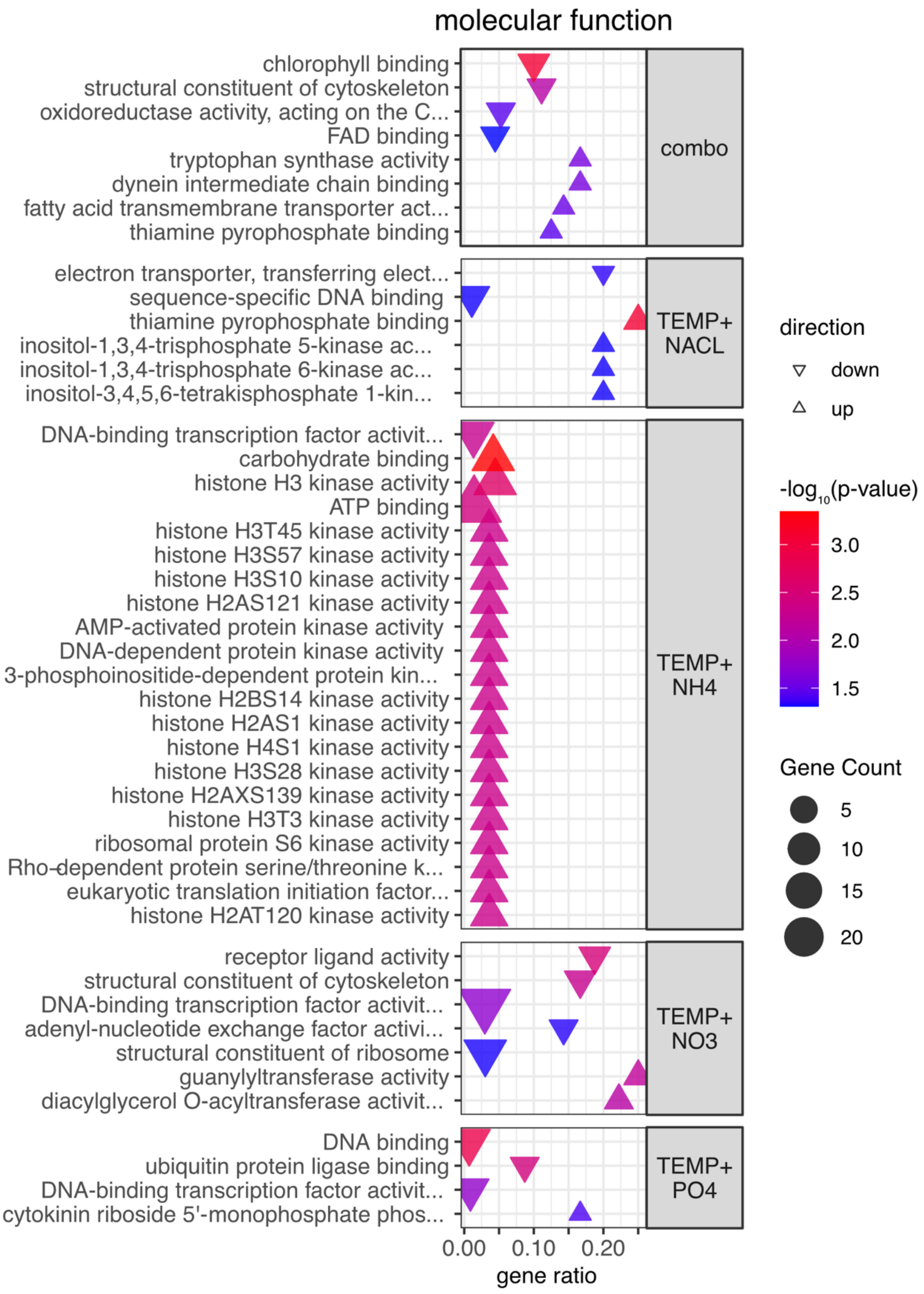


**Supplemental Figure 6: Molecular function GO terms suggest kinase signaling and DNA binding play roles in response to combinatorial treatments.** Significant

GO terms for molecular function in DEGs shared across combinatorial treatments (combo) and DEGs unique to specific combinations (y-axis). Gene ratio (x-axis) indicates the proportion of the gene list enriched for a GO term. Shape size indicates total number of genes enriched for a term; upward arrow indicates GO term significance among upregulated genes while downward arrow indicates enrichment in down regulated genes. Shape color corresponds to the degree of significance after Bonferroni correction for multiple tests. Kinase activity appears to be highly upregulated in genes unique to TEMP+NH4 and TEMP+NACL. In particular, upregulated DEGS in TEMP+NH4 are enriched for histone kinases. Among downregulated DEGs enrichment for GO terms related to DNA binding activity are found across several conditions including TEMP+NACL, TEMP+NH4, TEMP+NO3 and TEMP+PO4 conditions.

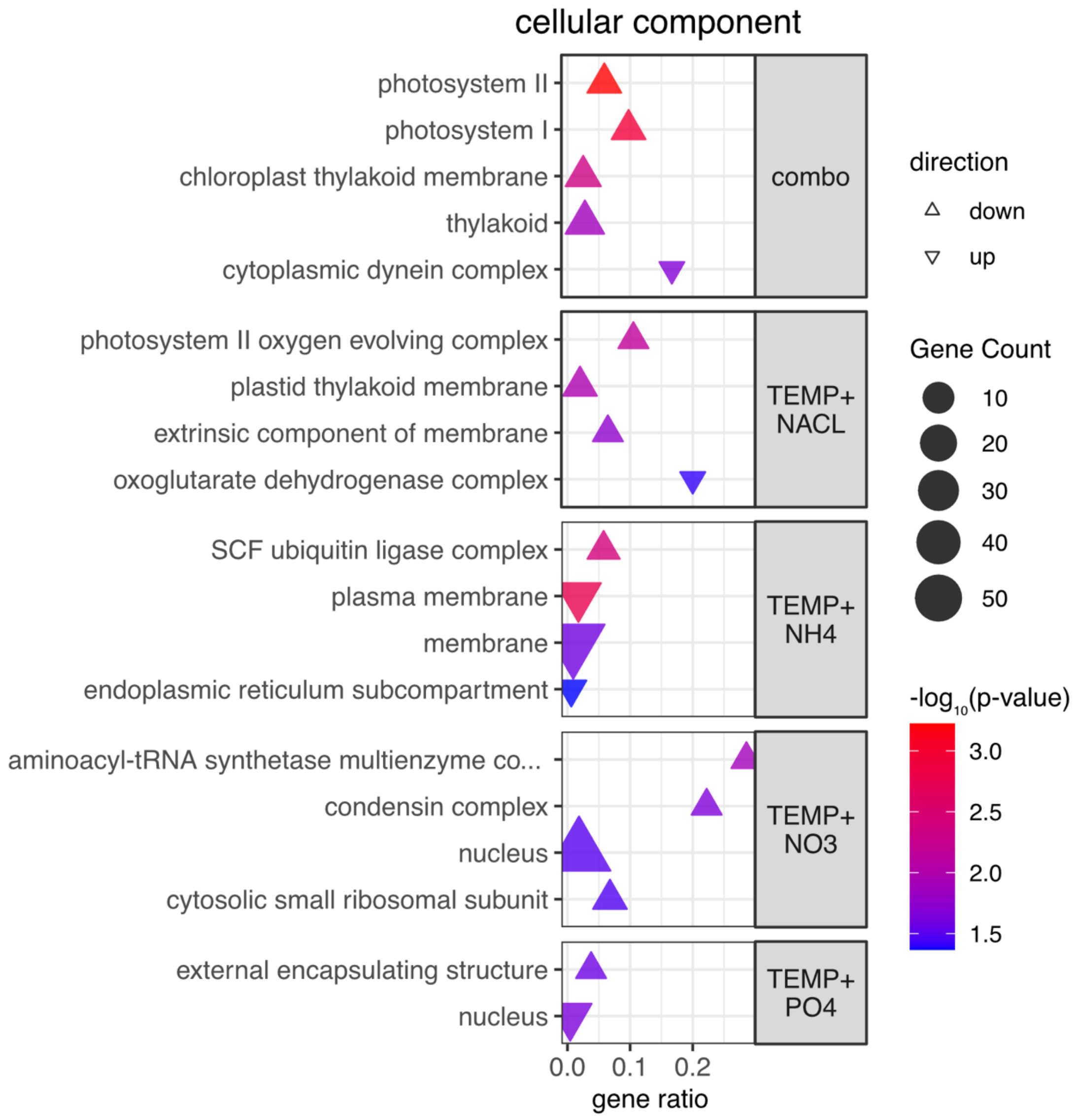


**Supplemental Figure 7: GO enrichments for Cellular compartment suggest changes in photosynthesis and protein production.** Significant GO terms for cellular compartment in DEGs shared across combinatorial challenge (combo) and DEGs unique to specific combinations (y-axis). Gene ratio (x-axis) indicates the proportion of the gene list enriched for a GO term. Shape size indicates total number of genes enriched for a term; upward arrow indicates GO term significance among upregulated genes while downward arrow indicates enrichment in down regulated genes. Shape color corresponds with the degree of significance after Bonferroni correction for multiple tests. Down regulation in genes associated with photosystem I, II and thylakoids suggest changes in photosynthetic processes. Similarly, down regulation of genes associated with terms related to protein production were enriched in TEMP+NO3 and TEMP+PH conditions.

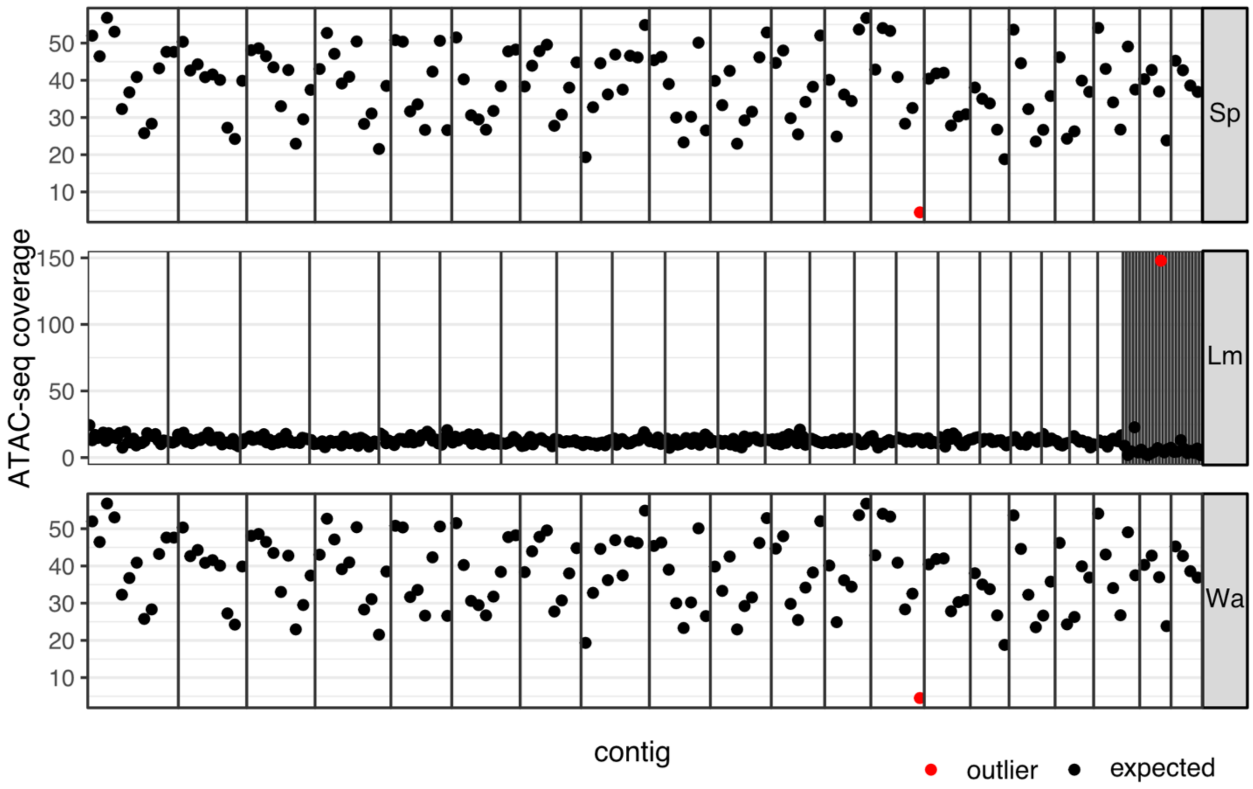


**Supplemental Figure 8: Coverage of ATAC-seq reads across duckweed genomes.** ATAC-seq read coverage (y-axis) over 1-Mb windows of the genome (x-axis) of 3 duckweed species; mitochondria and chloroplast genomes were excluded. Outliers (red points) were designated using the Z-score method with a threshold of $|Z| > 3$. The majority of read coverage across genomes scaffolds (vertical lines) falls within the expected range of coverage (black points).

A)

| rep | species | mapped reads | % nuclear mapping | peaks | friP | friT |
|---|---|---|---|---|---|---|
| 1 | Sp | 43491349 | 92 | 25524 | 0.52 | 0.30 |
| 2 | Sp | 26649820 | 77 | 25493 | 0.43 | 0.23 |
| 3 | Sp | 23000494 | 77 | 25702 | 0.43 | 0.23 |
| 4 | Sp | 21537560 | 88 | 25026 | 0.46 | 0.24 |
| 1 | Lm | 35359119 | 67 | 26098 | 0.32 | 0.20 |
| 2 | Lm | 27235057 | 78 | 25964 | 0.37 | 0.24 |
| 3 | Lm | 23569491 | 78 | 26561 | 0.32 | 0.20 |
| 1 | Wa | 10599472 | 94 | 29419 | 0.42 | 0.18 |
| 2 | Wa | 6384784 | 86 | 29458 | 0.44 | 0.17 |
| 3 | Wa | 5932739 | 86 | 30971 | 0.42 | 0.17 |

B)

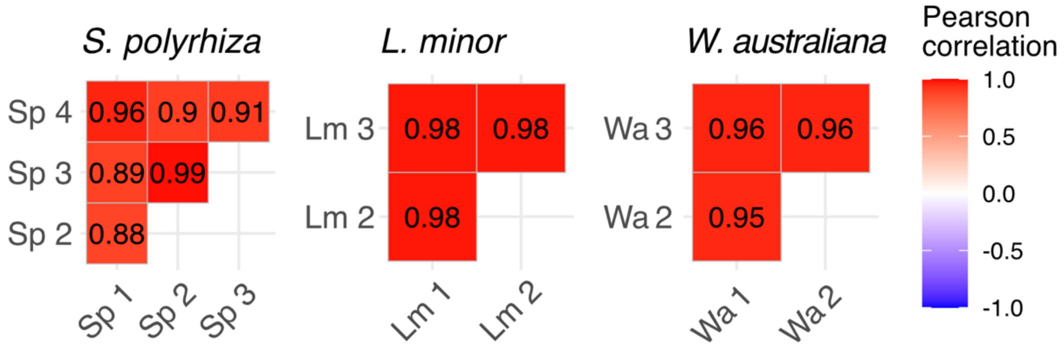


C)

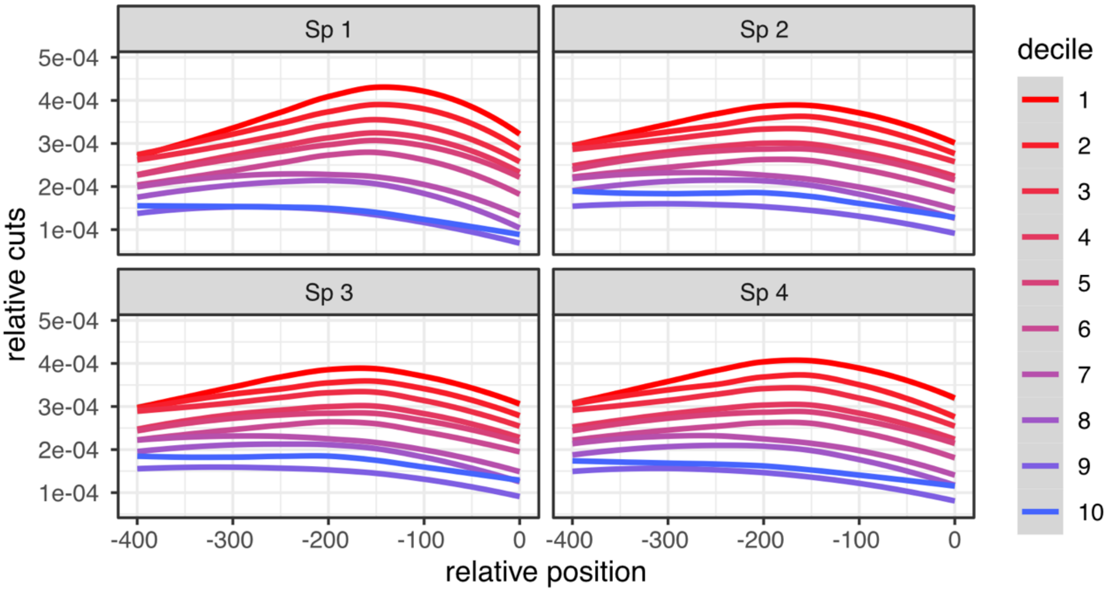

**Supplemental Figure 9: Chromatin accessibility and sample correlations suggest high-quality and replicable data.** A) Quality metrics for ATAC-seq samples in three duckweed species. The proportion of uniquely mapping reads that do not map to the chloroplast or mitochondrial genome (% nuclear mapping) was above 60% for all samples. Total peaks called using heuristic peak caller were similar across samples. Fraction of reads within peaks (FriP score is above 30% for all samples) and fraction of reads in the 400bp upstream of the TSS (FriT score) was above 20% in *S. polyrhiza* and *L. minor* and above 15% for *W. australiana*. B) Pearson replicate correlations, using the number of inserts falling into ACRs calculated from merged reads. C) Expression stratified accessibility plots of relative Tn5 insertions i.e. relative cuts (y-axis), calculated as the $cuts_{postion}$ / $cuts_{total}$, within the 400 bp upstream of the transcriptional start site (x-axis) for *S. polyrhiza* samples. Genes are stratified into expression deciles with 1 being the most expressed genes (red) and 10 being the least expressed (blue); genes with no detectable expression were not included.

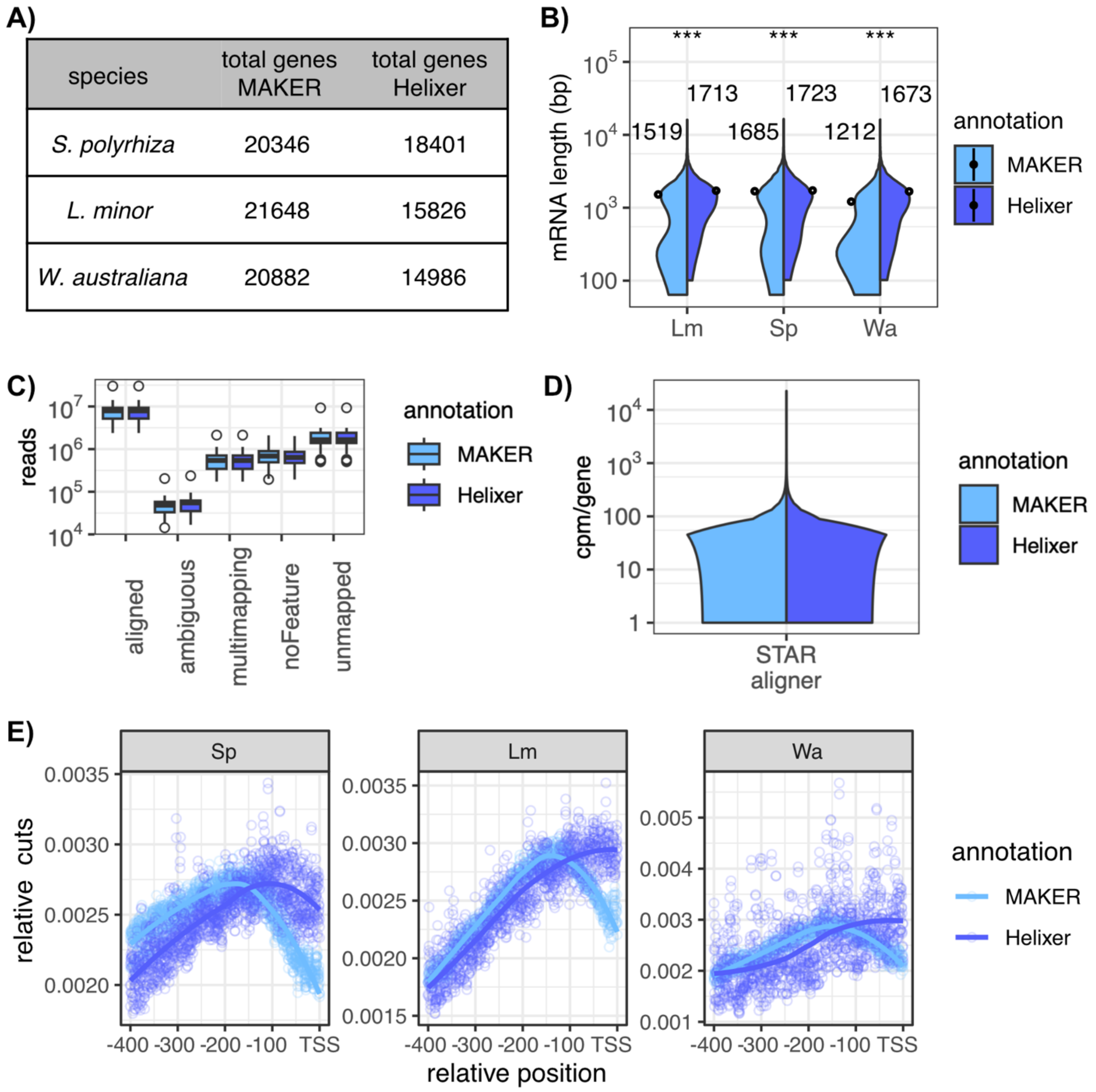


**Supplemental Figure 10: MAKER and Helixer annotations are of comparable quality.** A) Total number of annotated genes for three species of duckweed using two annotation methods: MAKER (expression-tag informed) or Helixer (deep neural net with Hidden Markov model). B) The distribution of the mRNA lengths between annotations is significantly different in each species (Kolmogorov-Smirnov test: *L. minor* <0.0005, *S. polyrhiza* <0.0005, *W. australiana*<0.0005) C) *S. polyrhiza* RNA-seq reads mapped using either MAKER or Helixer annotations do not show significant differences in the total number of reads that where uniquely mappable (aligned), mapping to ambiguous regions (ambiguous), mapping to multiple genes (multimapping), reads aligned to the genome but not to an annotated gene (noFeature) and reads that could not be mapped (unmapped). D) There was no significant difference between the distribution of reads per gene (y-axis) between MAKER annotation and the Helixer annotations. Genes showing CPM less than one excluded. E) The relative number of Tn5 inserts i.e. cuts (y-axis), calculated as the $cuts_{postion}$ / $cuts_{total}$, for the 400 bp region upstream of the TSS (x-axis). Points represent calculated relative cuts while solid lines represent LOESS-smoothed trends.

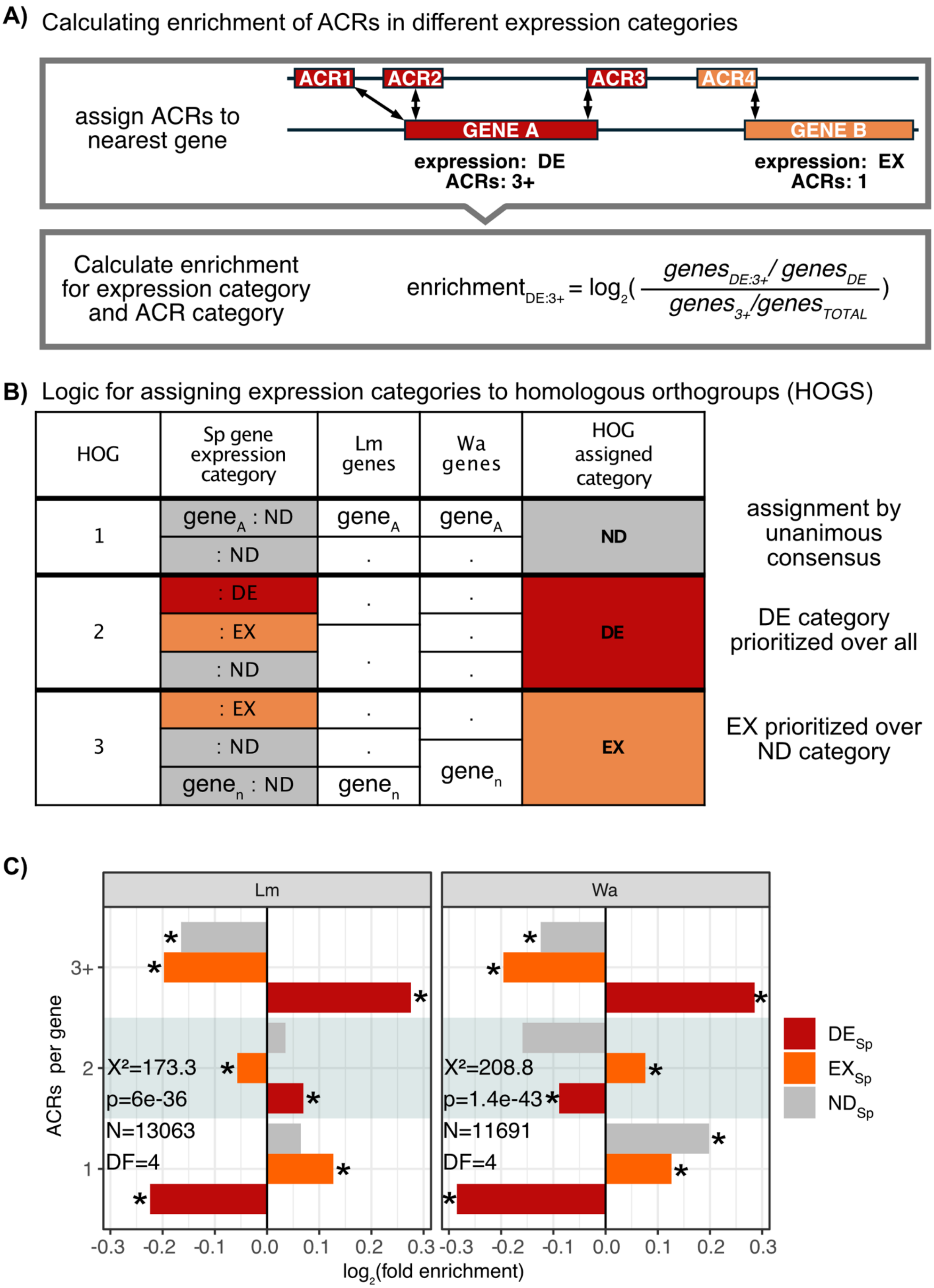
A) Calculating enrichment of ACRs in different expression categories
assign ACRs to nearest gene
ACR1
ACR2
ACR3
ACR4
GENE A
GENE B
expression: DE
ACRs: 3+
expression: EX
ACRs: 1
Calculate enrichment for expression category and ACR category
$enrichment_{DE:3+} = \log_2\left(\frac{genes_{DE:3+} / genes_{DE}}{genes_{3+} / genes_{TOTAL}}\right)$
B) Logic for assigning expression categories to homologous orthogroups (HOGS)
HOG
Sp gene expression category
Lm genes
Wa genes
HOG assigned category
1
$gene_A$ : ND
: ND
$gene_A$
$gene_A$
ND
2
: DE
: EX
: ND
DE
3
: EX
: ND
$gene_n$ : ND
$gene_n$
$gene_n$
EX
assignment by unanimous consensus
DE category prioritized over all
EX prioritized over ND category
C)
Lm
Wa
ACRs per gene
3+
2
1
X²=173.3
p=6e-36
N=13063
DF=4
X²=208.8
p=1.4e-43
N=11691
DF=4
-0.3 -0.2 -0.1 0.0 0.1 0.2 0.3
$\log_2$(fold enrichment)
$DE_{Sp}$
$EX_{Sp}$
$ND_{Sp}$

**Supplemental Figure 11: *L. minor* and *W. australiana* show enrichment of ACRs in genes that share orthology with *S. polyrhiza* environmentally responsive genes.** A) Top: representation of steps used to assign ACRs an expression category (DE, EX or ND) and to assign genes an ACR category (1, 2 or 3+). Bottom: calculation of enrichment score. B) Illustration of logic used to assign expression categories from *S. polyrhiza* measurements to *L. minor* and *W. australiana* genes based on homologous orthogroup clusters. C) Log2 fold-enrichment (x-axis) of *L. minor* and *W. australiana* genes showing different numbers of associated ACRs (y-axis). Expression categories are assigned based on orthology to genes in *S. polyrhiza*. Most prominent is the enrichment of ACRs near genes that share orthology to environmentally responsive genes in *S. polyrhiza* (red).

A)

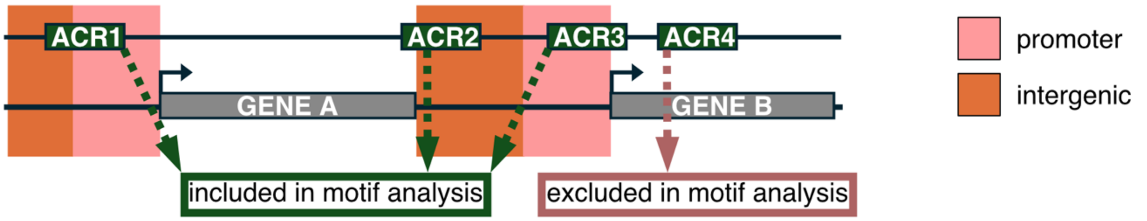


B)

| subset | ACRs |
|---|---|
| background | 12835 |
| temp up | 1401 |
| media up | 741 |
| combo up | 1815 |
| temp down | 1786 |
| media down | 282 |
| MFS down | 1808 |

C)

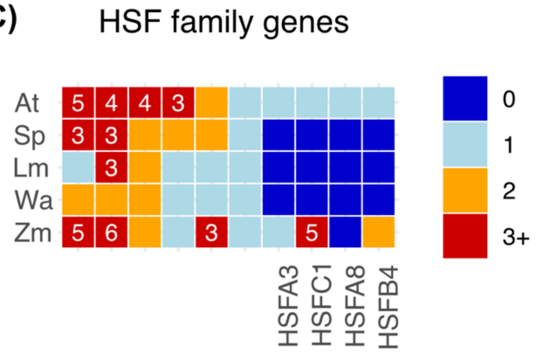


**Supplemental Figure 12: Sub-setting ACRs for motif enrichment analysis.** A) Only ACRs overlapping TSS-proximal or intergenic regions were used for motif analysis. B) Total ACR sequences used for DEGs from individual and combinatorial treatments. C) Heat shock factor family proteins are reduced in duckweed but still present.

Supplemental Tables:

| combination | deviation from additive effect | standard error | degrees of freedom | t-statistic | p-value |
|---|---|---|---|---|---|
| TEMP+HCL | 0.00681736 | 0.0296275 | 64 | 0.2301022 | 0.81874622 |
| TEMP+NACL | 0.045142 | 0.02460277 | 64 | 1.8348341 | 0.07117843 |
| TEMP+NH4 | 0.02754969 | 0.03351074 | 64 | 0.8221152 | 0.41406374 |
| TEMP+NO3 | -0.0547158 | 0.03182339 | 64 | -1.7193586 | 0.09038247 |
| TEMP+PO4 | 0.01279798 | 0.036496 | 64 | 0.3506679 | 0.72698914 |

**Supplemental Table 1: Growth rate in response to combinatorial treatments did not significantly deviate from the additive effect of corresponding individual treatments.**

| combination | observed DEGs | total genes | expected proportion | p-value |
|---|---|---|---|---|
| TEMP+HCL | 2872 | 15121 | 0.1856359 | 0.89 |
| TEMP+NACL | 3276 | 15121 | 0.1920508 | 8.63E-14 |
| TEMP+NH4 | 3152 | 15121 | 0.2247206 | 8.63E-06 |
| TEMP+NO3 | 3990 | 15121 | 0.1874876 | 3.71E-127 |
| TEMP+PO4 | 3093 | 15121 | 0.1852391 | 5.24E-09 |

**Supplemental Table 2: Combinatorial stress treatments showed significantly higher numbers of DEGs than expected from additive expectations of individual treatments**.

| species | ACR total | ACR proximal | ACR neighbor | ACR distal |
|---|---|---|---|---|
| AT | 34661 | 11531 | 13374 | 1944 |
| $SP_{7498}$ | 23472 | 3996 | 5984 | 1590 |
| SP | 27466 | 7967 | 9950 | 1656 |
| LM | 29124 | 8938 | 8760 | 2694 |
| WA | 33823 | 8129 | 10742 | 6684 |
| ZM | 35932 | 7032 | 11373 | 13124 |

**Supplemental Table 3: Similar numbers of ACRs and distance from gene distributions between species.** Comparison between species of total ACRs, ACRs overlapping the 400bp upstream of the TSS (proximal), ACRs closer than 2kb neighboring, and ACRs further that 2kb (distal).
